# A branching cell-fate decision in biofilm dispersal enables long-term surface persistence

**DOI:** 10.64898/2026.04.24.720661

**Authors:** Sandhya Kasivisweswaran, Jojo A. Prentice, Andrew A. Bridges

**Affiliations:** Department of Biological Sciences, Carnegie Mellon University; Pittsburgh, PA 15213, USA

## Abstract

Biofilms are the most ancient multicellular communities on Earth, representing a primitive developmental system that protects microbes from threats. Biofilm dispersal, whereby bacteria exit biofilms, is critical for the spread of pathogens to new infection sites. Here, using *Vibrio cholerae*, we show that dispersal events are accompanied by a branching cell-fate decision. While ∼90% of cells disperse, a viable subpopulation remains within a residual matrix. This post-dispersal biofilm community (PDBC) is established by the matrix protein RbmA and adopts a specialized anabolic program that enhances tolerance to antibiotics and bacteriophages. Our findings reveal that PDBCs act as a resilient “seed-bank” capable of rapidly re-populating the niche without requiring de novo matrix biosynthesis, providing a mechanistic basis for the recurrence and spread of chronic infections.

## Introduction

Bacteria commonly exist in surface-attached multicellular communities known as biofilms, a lifestyle that is thought to have arisen many times independently in the natural history of prokaryotes (*1*). The benefits of the biofilm state for resident bacterial cells include steric separation from environmental threats, stable surface colonization, and enhanced opportunities for metabolic cooperation and gene transfer (*2–5*). Biofilm formation occurs through the secretion of adhesive extracellular matrix components, such as polysaccharides, adhesin proteins, and extracellular DNA, which drive cell-cell and cell-surface interactions (*6–8*). Due to the greater stress tolerance and inherent difficulty associated with removing biofilms from surfaces, biofilm communities are notorious drivers of hospital-acquired infections and biofouling in many industries (*9*, *10*).

Despite the advantages conferred to resident bacterial cells, existence in densely packed biofilm communities can impose nutrient limitation and restricts dissemination (*11*). Thus, bacteria face a tradeoff between a vulnerable exploratory lifestyle (the planktonic state) and a sessile, crowded arrangement that offers protection (the biofilm state). To navigate this tradeoff, bacteria have evolved elaborate regulatory networks that use cellular and environmental signals to inform transitions between the biofilm and free-swimming lifestyles (*12*, *13*). This is exemplified by the ability of many bacteria to actively disassemble their biofilms from within (referred to as dispersal or dispersion), facilitating the spread of liberated bacteria to new environmental locations or infection sites for colonization (*14*). In the case of global pathogen *Vibrio cholerae*, a model organism for the biofilm lifecycle and the causative agent of cholera disease, dispersal is thought to facilitate host intestinal colonization and is therefore critical for disease transmission (*15*, *16*). While the molecular mechanisms of biofilm assembly have been well-characterized in numerous bacterial species, the process of biofilm dispersal remains comparatively understudied (*17*). Biofilm dispersal is often actively regulated, and can be conceptualized as involving the following mechanistic steps: (1) detection of signals that promote dispersal, (2) cessation of extracellular matrix production, and (3) weakening of adhesive interactions and/or the initiation of motility to facilitate cell exit (*14*, *18*). Biofilm dispersal is typically portrayed as the terminal phase of the biofilm lifecycle, leaving the fate of the residual biofilm communities after dispersal largely unexplored. Post-dispersal structures could support persistent surface colonization, potentially leading to chronic infection or erosion, making their study essential.

Much like tissues in higher organisms, biofilm cells are known to exhibit phenotypic heterogeneity, driven by inherent gene expression noise, stochastic partitioning of proteins at cell division, and metabolic or signaling gradients that develop in spatially structured communities (*11*, *19–23*). Yet, how this physiological heterogeneity manifests in biofilm dispersal remains unclear. In *V. cholerae,* the dispersal process is regulated by the integration of quorum sensing and stress-responsive regulatory pathways, linking population-level cues with local environmental conditions (*24*, *25*). High-resolution microscopy of *V. cholerae* and *Pseudomonas aeruginosa* biofilms has revealed that dispersal is spatially patterned and heterogeneous, leaving behind a subset of cells after its completion, suggesting that biofilm dispersal could represent a point of divergence of cell fates (*26–28*). However, whether residual cells represent dispersal by-products, or a specialized population with ecologically distinct functions remains unknown. Together, these findings raise fundamental questions about how dispersal outcomes might vary at the single-cell level, how phenotypic heterogeneity biases which cells disperse versus remain, and what the long-term fate of the post-dispersal biofilm community is -- specifically whether residual biofilm structures can recover, persist, or even re-seed future biofilm growth.

Here, we uncover the molecular drivers and ecological consequences of phenotypic differentiation during *V. cholerae* biofilm dispersal. Using single-cell, high-resolution confocal microscopy with far-red fluorogenic probes, we reveal that biofilm dispersal reproducibly leaves behind a viable, matrix-encased subpopulation of cells that persists as a structured post-dispersal biofilm community (herein referred to as PDBC). We identify the matrix protein RbmA as necessary and sufficient for PDBC establishment. Transcriptomic profiling reveals that cells remaining in PDBCs exhibit a distinct physiology relative to their isogenic dispersed counterparts, suggesting that they play divergent ecological roles. PDBCs exclude non-resident bacteria, tolerate diverse environmental stresses, and retain the capacity to rapidly regenerate biofilms without synthesis of new biofilm matrix. Collectively, our findings suggest that dispersal is not a uniform endpoint for the biofilm lifecycle; phenotypic heterogeneity generates an exploratory planktonic fraction and a resilient post-dispersal biofilm community that functions as a seed-bank, persisting on surfaces and supporting the population in fluctuating environments, consistent with a bet-hedging strategy.

### *V. cholerae* biofilm cells exhibit phenotypic diversity during dispersal

Phenotypically flexible *V. cholerae* strains undergo a complete biofilm lifecycle when inoculated in static growth conditions: after initial founder cell attachment, clonal microcolony biofilms develop for ∼9 hours, after which coordinated dispersal occurs as quorum and starvation signals accumulate (*24*, *25*, *29*). To assess *V. cholerae* biofilm dispersal at high resolution, we performed confocal microscopy of this lifecycle using cells labeled with the genetically encoded far-red fluorogen binding protein dL5 (hereafter referred to as dL5-red) (*30*). Consistent with prior work, we observed that a subpopulation of cells remained biofilm-associated after dispersal reached completion (the point when no further cell departures occurred) (Fig. 1A, B, Movie S1) (*26*). We found that, at this point, 10 ± 4% (Mean ± SD) of cells remained in biofilms while the bulk of the population had transitioned to an exploratory state (Fig. 1C). Examination of PDBCs at substantially later timepoints confirmed that these cells remained surface associated even on longer timescales of up to 48 hours post-inoculation (Fig. S1A). Notably, peak biofilm size at the onset of dispersal had only a modest correlation (r = -0.578) with the percentage of cells remaining post-dispersal (Fig. 1C, Fig. S2), suggesting that dispersal-associated phenotypic differentiation does not strongly depend on peak biofilm size. Thus, biofilm dispersal in *V. cholerae* appears to be accompanied by a cell-fate divergence, generating two phenotypically distinct cell populations.

To investigate why a subpopulation of cells remain biofilm-associated after the completion of dispersal, we examined two possibilities – (i) that remaining cells are dead, or (ii) that remaining cells are alive but trapped within a partially degraded biofilm matrix. To this end, we first investigated the viability of non-dispersed cells by staining PDBCs with SYTOX Green, a membrane-impermeable nucleic acid dye, to selectively label dead cells with a compromised cell membrane. We observed SYTOX staining in only 1-5 cells per PDBC, confirming that the vast majority of cells remaining in PDBCs were alive (Fig. 1D, Fig. S1B). Given that biofilm-associated cells have previously been suggested to enter a viable but non-culturable (VBNC) state, we sought to test the ability of PDBC cells to seed culture growth (*31*). To do so, we separated dL5-red-expressing dispersed cells from non-dispersed cells by pipetting, washed equal numbers of each population into fresh growth media, and assessed culture growth through bulk fluorescence signal. We observed that both dispersed and PDBC subpopulations were culturable and grew equally well when fresh nutrients were provided (Fig. 1E). Together, these results demonstrate the presence of a viable, culturable subpopulation of cells that remain entrapped in *V. cholerae* biofilms after dispersal. We propose that such phenotypic variation in biofilm dispersal is an emergent property of biofilm matrix architecture and could constitute a bet-hedging strategy, that confers a survival advantage on *V. cholerae* in uncertain environments (*32*).

**Figure 1:**
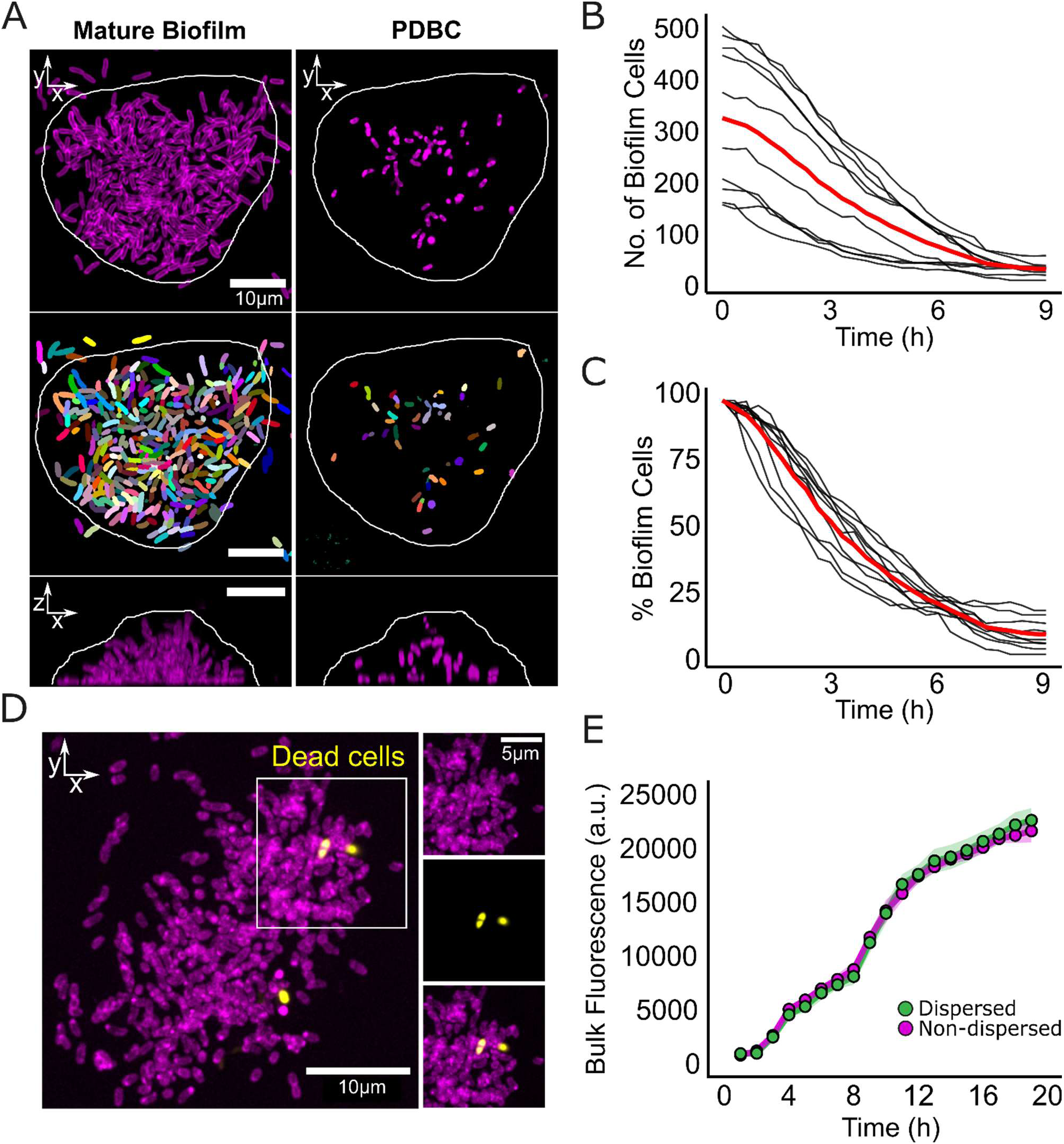
A viable population of *V. cholerae* cells remains surface-attached after the completion of biofilm dispersal. (A) Representative images of wild-type *V. cholerae* biofilms before (left panels) and after biofilm dispersal (right panels) at 10 and 16 hours post-inoculation respectively. Confocal micrographs of a single bottom slice (top panels), their segmentation into individual cells (false-colored) (middle panels) and x-z projections of biofilms (bottom panels). Cells constitutively expressed dL5-red and are displayed in magenta. White line indicates biofilm boundary. (B) Number of biofilm-associated cells in 10 replicate biofilms over the course of dispersal as measured from single-cell segmentation of spinning-disk confocal sections imaged at 20-min intervals for 9 hours. Timepoint zero represents where biofilm biomass is at a peak. Red line represents the mean of all replicates. (C) Data in B represented as a percentage of the number of cells in a mature biofilm over the course of dispersal. (D) Representative sum projection of a PDBC stained with the nucleic acid stain SYTOX Green, *N* = 3 biological replicates (Fig. S1B). Magenta represents all cells constitutively expressing dL5-red, and yellow represents SYTOX signal. (E) Bulk fluorescence intensity of dispersed and PDBC cell populations when grown separately in fresh growth media at equal starting cell numbers. Cells of both populations expressed dL5-red and fluorescence output was measured in bulk culture using a plate reader. Points represent means of *N* = 3 biological and 3 technical replicates of each strain, ± SD (shaded). a.u., arbitrary units.

### Remaining matrix components provide structural integrity to PDBCs

Given the importance of extracellular matrix components in establishing the biofilm lifestyle, we wondered whether specific biofilm structural components were also responsible for retaining cells in the PDBC. Timelapse imaging of biofilm dispersal showed that dispersed cells exhibited vigorous motility but were unable to re-enter the biofilm regions from which they had dispersed (Movie S1). Indeed, when we performed a two-color imaging experiment in which we mixed mNeonGreen-labeled dispersed cells with dL5-red-labeled PDBCs, dispersed cells were unable to invade PDBCs (Fig. 2A, Movie S2). The exclusion of external cells suggests that the biofilm matrix, which is comprised of vibrio polysaccharide (VPS) and matrix proteins, may remain intact after dispersal and provide structural integrity to PDBCs. We reasoned that remaining matrix components could simultaneously trap the non-dispersing population and exclude external cells. To examine this possibility, we probed the localization and abundance of polysaccharide and protein matrix components in PDBCs. Indeed, the VPS matrix was abundant in post-dispersal biofilms, both immediately surrounding remaining cells and in biofilm regions from which cells had successfully dispersed (Fig. 2B, Fig. S3A). Given that biofilm scaffolding protein RbmA was previously shown to be essential for excluding invading cells from mature biofilms, we hypothesized that it might play a similar role in PDBCs (*33–35*). Indeed, much like the VPS matrix, we observed significant RbmA-3xFLAG staining in PDBCs (Fig. 2C, Fig. S3B), both immediately surrounding PDBC cells as well as in regions from which cells had successfully dispersed. Together, these data show that PDBCs retain structural matrix components including VPS and RbmA, suggesting that dispersal does not involve complete enzymatic digestion of the biofilm matrix. This preserved matrix limits external cell invasion and creates a distinct ecological niche for resident cells.

**Figure 2:**
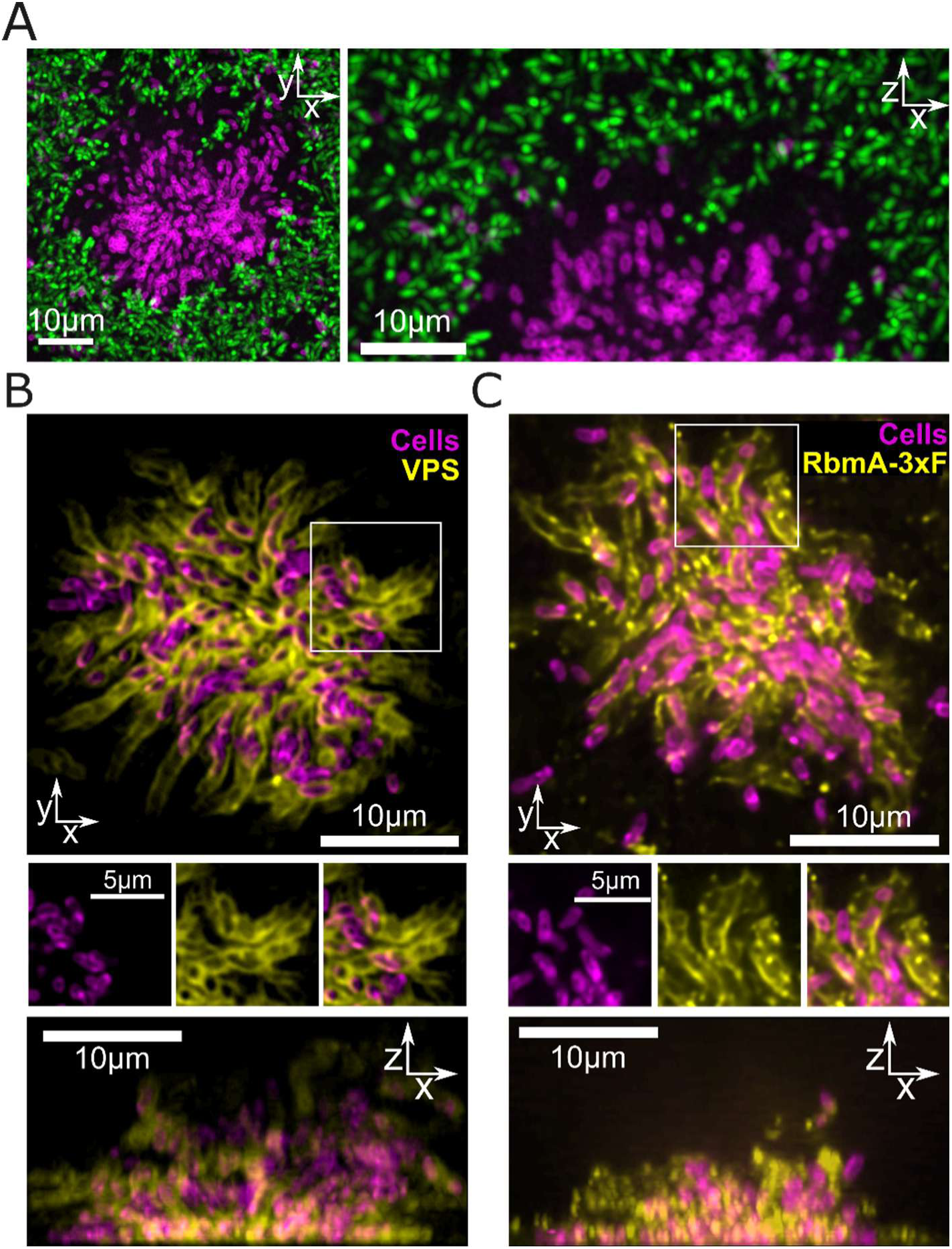
Biofilm matrix components persist in PDBCs. (A) Representative confocal micrographs of a wild-type *V. cholerae* PDBC constitutively expressing dL5-red (magenta) with dispersed cells expressing mNeonGreen (green). Time projections of representative x-y (left) and x-z (right) planes. (B) Representative confocal micrographs of the bottom slice (top panel), magnified view of the boxed region (middle panel), and x-z projection (bottom panel) of a post-dispersal biofilm, probed for VPS using 1 μM GFP-conjugated Bap1 protein (*36*). Magenta represents cells constitutively expressing dL5-red, and yellow represents Bap1-GFP fluorescence. (C) Representative confocal micrographs of the bottom slice (top panel), magnified view of the boxed region (middle panel), and x-z projection (bottom panel) of a post-dispersal biofilm, probed for C-terminal tagged RbmA-3xFLAG using an anti-FLAG-M2-Cy3 antibody. Magenta represents cells constitutively expressing dL5-red, yellow represents anti-FLAG-antibody fluorescence. *N* = 3 biological replicates for all images displayed.

### RbmA is required to establish the PDBC fate in *V. cholerae*

The preservation of matrix integrity in PDBCs suggests that remaining adhesive components could retain the non-dispersing population. We previously observed that the Δ*rbmA* mutant strain forms biofilms which disperse to completion, with no cells remaining at the final timepoint (replicated in Fig. 3A, Movie S3), indicating that RbmA is required for retaining the PDBC cell population (*26*). To assess whether RbmA is sufficient to drive the PDBC cell fate, we increased RbmA levels beyond that of the wild-type strain using an arabinose-inducible promoter and found that overexpression of rbmA led to reduced cell departures without impacting overall peak biofilm biomass (Fig. 3B, S4). Having established the link between RbmA levels and the retention of cells in PDBCs, we examined the temporal changes in *rbmA* gene expression throughout the biofilm lifecycle (growth and dispersal). To this end, we first generated a fluorescent transcriptional reporter by fusing the promoter of *rbmA* (*P_rbmA_*) to *dL5-red*. Quantification of *P_rbmA_-dL5-red* fluorescence signal relative to a constitutive reporter (*P_tac_-AM2-2*) from whole biofilms over time revealed that *rbmA* expression peaked before biofilm biomass peaked, after which promoter activity steadily decreased throughout biofilm dispersal (Fig. 3C). Given our observation of robust RbmA-3xFLAG protein staining in PDBCs (Fig. 2C), we next investigated how RbmA-3xFLAG protein levels changed over the course of the biofilm lifecycle. Similar to the promoter fusion, RbmA-3xFLAG staining peaked before biofilm biomass peaked, however, unlike the gene expression reporter, RbmA-3xFLAG persisted throughout the dispersal process (Fig. 3D, E, Fig. S5, Movie S3). This demonstrates that *rbmA* gene expression drives the accumulation of RbmA in developing biofilms and that, despite transcriptional downregulation preceding the onset of dispersal, the RbmA protein persists throughout dispersal. To evaluate radial trends in RbmA localization throughout biofilm development, we quantified RbmA-3xFLAG signal in radial bins moving outward from the biofilm core on the substrate (Fig. 3F and Fig. S6). Notably, when biofilm biomass peaked, RbmA-3xFLAG localization was highest at the biofilm core (Fig. 3D, F). We note that this localization is consistent with the majority of non-dispersed cells remaining in the core of the PDBC (*26*). Taken together, these findings suggest that RbmA accumulation during biofilm development and its persistence through dispersal help maintain post-dispersal matrix structure that retains a subpopulation of cells in the PDBC.

**Figure 3:**
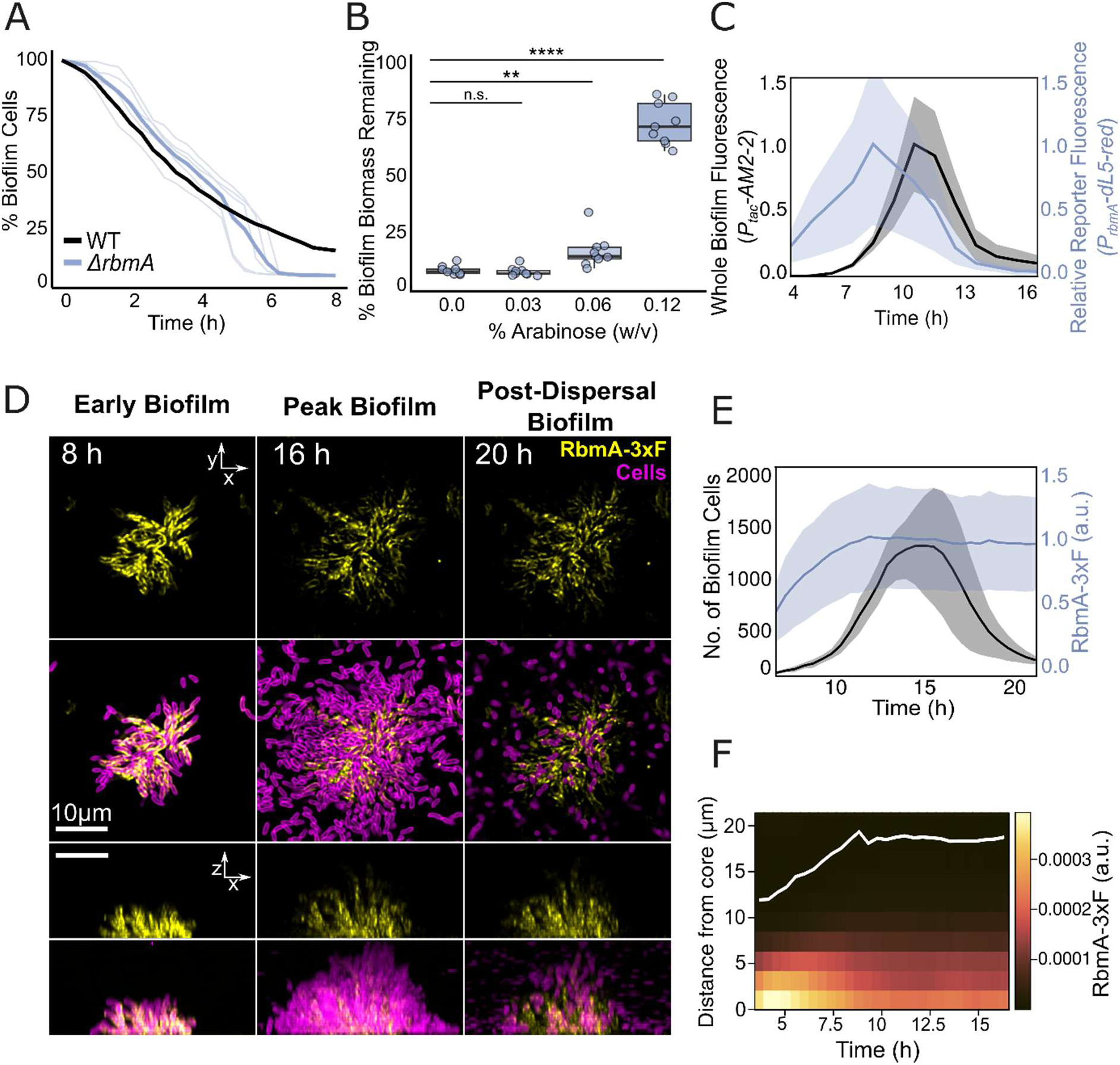
RbmA is necessary and sufficient to establish V. cholerae PDBCs. (A) Percentage of biofilm-associated cells relative to the number of cells in a mature biofilm over dispersal, for the indicated strains. Gray lines represent individual Δ*rbmA* replicates, blue line represents mean of Δ*rbmA* biofilms (*N* = 5 replicates), and black line represents the mean of wild-type biofilms (*N* = 10 replicates). Time point zero represents where biofilm biomass is at peak. (B) Box and whisker plot of the percentage of non-dispersed cells relative to the peak biofilm biomass in a Δ*rbmA* strain induced with the indicated arabinose concentrations (% w/v). Box plot represents interquartile range (box limits), mean, and maximum and minimum values (whiskers). Points represent all replicate values. *N* = 3 biological replicates and 3 technical replicates for each condition. Statistical analysis was performed using one-way ANOVA with Dunnett’s multiple comparisons test to compare each sample induced with arabinose to the uninduced sample. ****, *P* ≤ 0.0001; **, *P* > 0.01; ns, *P* > 0.05. ns., not significant. (C) Blue trace: *P_rbmA_–dL5-red* fluorescence over time measured from low-magnification spinning-disk confocal images (20× objective) of whole biofilms and normalized to a constitutive reporter (*P_tac_-AM2-2*) expressed in the same cells to yield relative promoter activity. Black trace: Whole biofilm *P_tac_-AM2-2* fluorescence plotted as a proxy for total biofilm biomass. Both traces are internally normalized to their respective maxima. *N* = 3 biological replicates. Lines represent means, ± SD (shaded). (D) Representative confocal micrographs of wild type V. cholerae biofilms expressing 3xFLAG-tagged RbmA through early (left panel), peak (middle panel), and post-dispersal (right panel) stages. Magenta represents cells constitutively expressing dL5-red, and yellow represents anti-FLAG antibody fluorescence. Top panels show x-y slices at the biofilm base (RbmA-3xFLAG only, and an overlay), and bottom panels show x-z projections (RbmA-3xFLAG only and an overlay) of the same biofilms. (E) Quantification of total fluorescence signal from RbmA-3xFLAG (blue line) in a biofilm, and number of biofilm cells (black line) through the course of biofilm development. Fluorescence signal is internally normalized to its maximum. Lines represent means of 5 replicate biofilms, ± SD (shaded). (F) Representative kymograph of a single biofilm, quantifying average RbmA-3xFLAG fluorescence intensity over time and at the indicated radial distances from the biofilm core. White line represents biofilm boundary (Fig. S6). a.u., arbitrary units.

### Two physiologically distinct cell populations exist after *V. cholerae* biofilm dispersal

Physical segregation of a clonal biofilm into dispersing and non-dispersing subpopulations raises the possibility that the two populations differ in gene expression and ecological function. To test this possibility, we compared the transcriptional profiles of dispersed and PDBC subpopulations. We reasoned that this approach could potentially reveal (1) gene expression programs underlying the establishment of the two cell states and (2) transcriptional divergence arising from existing in distinct physiological niches (i.e., free-swimming versus remaining within the PDBC matrix). The analysis revealed that dispersed and non-dispersed cells exist in markedly different transcriptional states with ∼20% of the transcriptome (765 genes) exhibiting significant differential expression (Fig. 4A, Table S1). Given that dispersed cells were highly motile while non-dispersed cells remained surface-associated and matrix-embedded, we reasoned that expression of known biofilm and/or motility factors could underlie these opposing phenotypes. However, we found that most genes in the *vps* matrix operons were not significantly differentially expressed in the two populations, nor were flagellar structural and biosynthesis genes (Fig. S7). Further, Gene Set Enrichment Analysis (GSEA) using known biofilm and motility genes confirmed that neither pathway, as a whole, was significantly differentially expressed across the two cell populations (enrichment *P-value* = 0.8 for both, Table S2). Consistent with this observation, measurements of intracellular levels of secondary messenger cyclic-di-GMP (c-di-GMP), a central regulator of motility and biofilm formation, did not significantly differ between the dispersed and PDBC cell populations (Fig. S8) (*37*, *38*). However, despite this pathway-level trend, a subset of biofilm-related genes, including *rbmA*, were modestly upregulated in the PDBC cells (Fig. S7). We reason that cell fate differentiation could be driven by transient gene expression programs that occur earlier in biofilm development.

**Figure 4:**
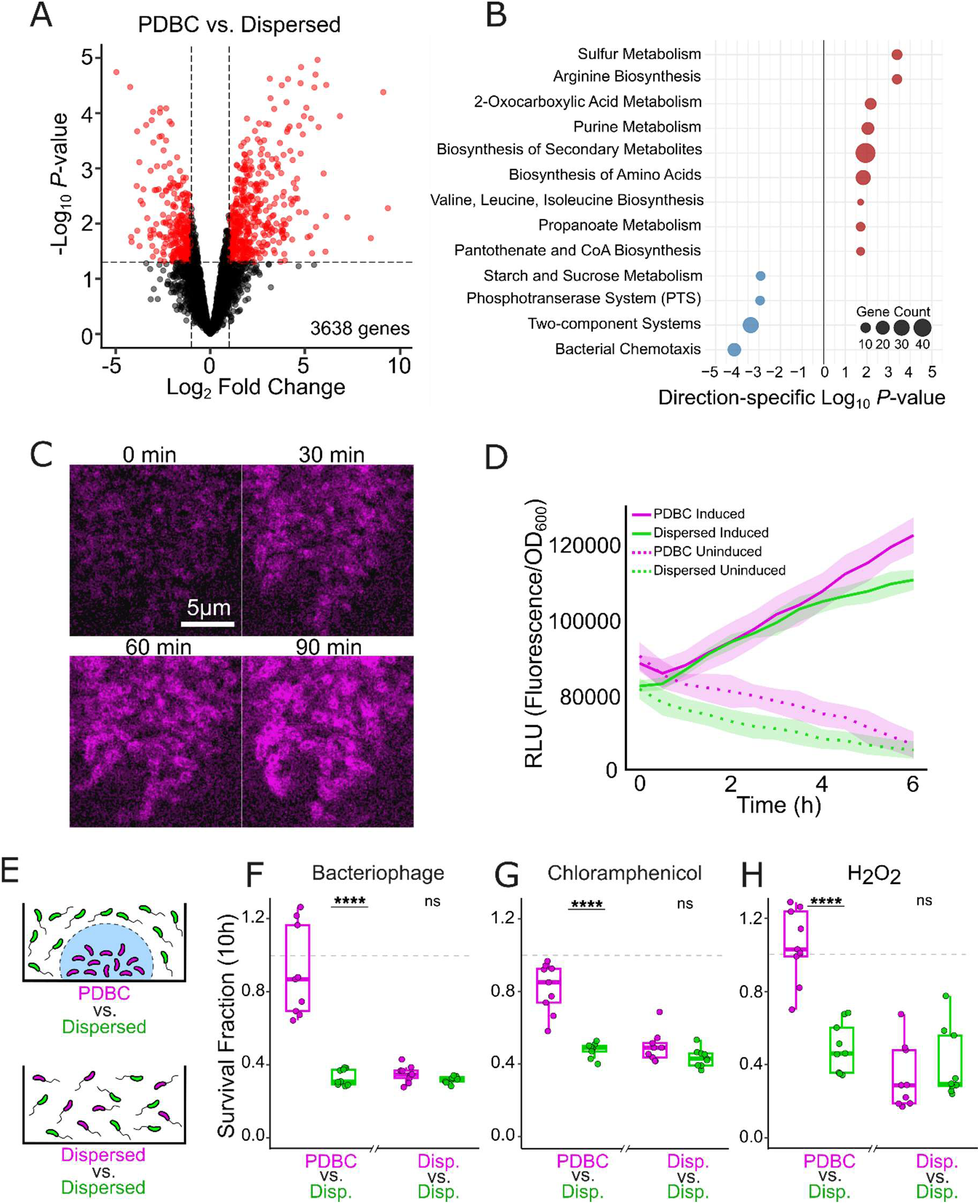
PDBC cells exhibit differentiated gene expression and survive challenges better than dispersed cells. (A) Volcano plot showing all quantified genes in PDBC cells compared to their dispersed counterparts. A Log_2_fold > 1 and *P*-value < 0.05 were used as significance cutoffs (dashed lines, red points are significant). *N* = 3 biological replicates. (B) Bubble plot representing KEGG pathway overrepresentation analysis. Elevated (red) and decreased (blue) pathways are for PDBC cells relative to dispersed cells. (C) Confocal micrographs (bottom slice) of cells in a PDBC after induction of *P_tac_-dL5-red* with 100 µM IPTG in a strain carrying LacI, at the indicated timepoints. Magenta represents dL5-red fluorescence, *N* = 3 biological replicates. (D) Quantification of bulk dL5-red fluorescence induction after the addition of 100 µM IPTG for the indicated cell populations. Relative Light Units (RLU) were calculated by normalizing fluorescence intensity by optical density (OD_600_) at each time point. Lines represent means of *N =* 3 biological and 3 technical replicates for each condition, ± SD (shaded). (E) Schematic of the two experimental conditions for competition assays in the presence of environmental threats. (F) Dispersed and PDBC cell population survival after 10 hours of competition in the presence of lytic bacteriophage S5 at MOI = 0.1. Survival fraction is normalized to untreated samples competed in the same conditions. To compare population sizes using fluorescence intensity of dL5-red and mNeonGreen, a fluorescence intensity to OD_600_ calibration was performed alongside each experiment in monoculture (see Methods for more information). Plot displays the mean (lines in the box), interquartile range (box limits), and maximum and minimum values within 1.5 times the interquartile range of the mean (whiskers). Dashed grey line is marked at a survival fraction of 1, representing no difference in survival upon treatment. Points represent all replicates. *N* = 3 biological replicates and 3 technical replicates for each condition. Two-sided unpaired *t* tests were performed for statistical analysis. ****, *P* ≤ 0.0001; ns, *P* > 0.05. ns., not significant. (G) As in F, in the presence of the translational inhibitor chloramphenicol at a sub-minimal inhibitory concentration of 3.1 µM. (H) As in F in the presence of 125 µM hydrogen peroxide. Disp., dispersed.

Examination of additional transcriptional differences between the two populations revealed that PDBC cells exhibited significant upregulation of core anabolic processes including biosynthesis of amino acids and purines, sulfur assimilation, Coenzyme A biosynthesis, and 2-oxocarboxylic acid metabolism (Fig. 4B). These results suggest that these cells are not metabolically quiescent but instead exhibit a robust anabolic program, allowing metabolism to continue in the PDBC. Consistent with this understanding, GSEA of stringent response genes, a nutrient limitation program associated with translational arrest and cellular dormancy, showed no significant enrichment (Table S2) (*39–41*). To experimentally assess whether PDBC cells are transcriptionally and translationally active, we generated an inducible, LacI-controlled *P_tac_-dL5-red* strain (see Methods for strain details). Upon induction, PDBC cells exhibited fluorescence above the background within 30 minutes, with similar induction kinetics to the dispersed population, demonstrating that these cells actively synthesize RNA and protein (Fig. 4C, D, Movie S5). We next examined the dispersed population, which, relative to the PDBC cells, displayed greater expression of pathways associated with carbon uptake and utilization, including the phosphotransferase system responsible for sugar import, as well as chemotaxis genes. Together, these findings suggest that PDBC cells adopt a transcriptional program characterized by increased anabolic processes, presumably to withstand the nutrient-limited conditions of the biofilm interior, while dispersed cells engage an exploratory program characterized by chemotaxis and nutrient scavenging. Taken together, cell fate differentiation is not distinguished by comprehensive differentiation in canonical biofilm or motility programs, but rather by metabolic specialization.

We reasoned that the differentiation of cells into dispersed and PDBC populations with distinct metabolic profiles could confer differing ecological roles to each phenotype. While most cells disperse from the biofilm to explore new territories, a subpopulation of cells stays behind, potentially to exploit the benefits of the remaining biofilm. Given that mature biofilm-associated cells are known to be shielded from threats such as antibiotics and bacteriophage infection, we wondered whether cells in the PDBC were afforded these same protective benefits (*4*, *42*). To test this possibility, we competed differentially fluorescently labeled dispersed cells and PDBC cells in the presence or absence of environmental stressors and measured the relative survival of each population using calibrated optical density for each fluorescence channel (see Methods, Fig. 4E, S9). Competing cell populations in the presence of lytic bacteriophage S5 revealed a ∼2.5-fold survival advantage for cells remaining in PDBCs relative to the dispersed population (Fig. 4F). Similarly, cells in the PDBC exhibited a ∼3-fold survival advantage in the presence of sub-MIC concentrations of the translational inhibitor chloramphenicol, and a ∼2-fold survival advantage during exposure to oxidative stress, which *V. cholerae* encounters both in infection and in environmental settings (Fig. 4G, H). Together, these data support a model where cells in the PDBC retain protective benefits of the biofilm lifestyle, representing a stress-tolerant niche for cells to tide over harsh environmental conditions, while their dispersed counterparts explore new territories.

### PDBC cells regenerate biofilms with minimal de novo matrix gene expression

The persistence of a stress-tolerant subpopulation of cells in the PDBC, that exclude invading competitors, suggests that the PDBC may act as a stable reservoir capable of re-populating an environmental niche when conditions improve. To examine this possibility, we assessed whether cells in the PDBC could regenerate biofilms when washed into fresh media following the completion of dispersal. Indeed, PDBCs regenerated mature biofilms of similar size to that of the initial biofilms (Fig. 5A-C). A subsequent dispersal event then occurred, again releasing the majority of cells into a free-swimming state, with a subpopulation remaining in the biofilm. This process could be repeated for at least four cycles (the most we tried), highlighting the capacity of non-dispersed cells to maintain population continuity under fluctuating environmental conditions (Fig. 5A, Movie S6).

**Figure 5:**
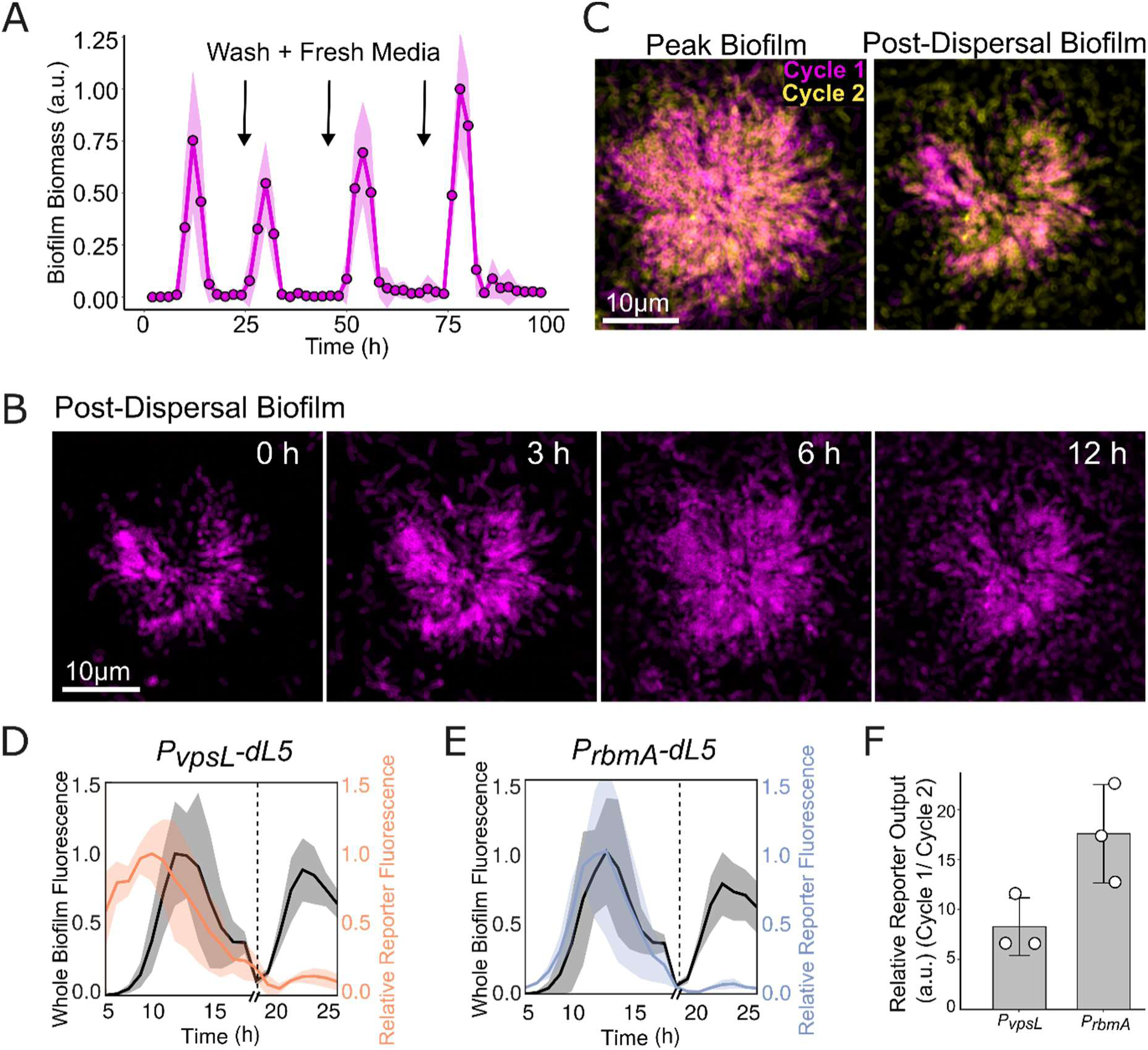
PDBCs reseed biofilms in fresh nutrient conditions. (A) Biofilm biomass measured using brightfield microscopy over four cycles of biofilm formation and dispersal, normalized to the peak biofilm biomass across the experiment. Black arrows indicate when washing was performed to remove dispersed cells and provide fresh growth media to PDBCs. Line represents the mean of *N* = 6 biological and 3 technical replicates each, ± SD (shaded). (B) Representative confocal micrographs (bottom slice) of a PDBC at indicated times after the addition of fresh growth media. (C) Representative confocal micrographs (bottom slice) of peak and post-dispersal biofilms during and after the first (magenta) and second (yellow) rounds of biofilm growth and dispersal. *N =* 3 biological replicates for all images displayed. (D) Orange trace: *P_vpsL_–dL5-red* fluorescence over time measured from low-magnification spinning-disk confocal images (20× objective) of whole biofilms and normalized to a constitutive reporter (*P_tac_-AM2-2*) expressed in the same cells to yield relative promoter activity. Black trace: Whole biofilm *P_tac_-AM2-2* fluorescence plotted as a proxy for total biofilm biomass. Dashed line indicates the time point where dispersed cells were removed and PDBCs were washed into fresh media. Both traces are internally normalized to their own maxima across both cycles. *N* = 3 biological replicates. Lines represent means, ± SD (shaded). (E) Same as in D for *rbmA.* Blue line represents relative *P_rbmA_–dL5-red* fluorescence over time. (F) Ratio of peak intensities from fluorescent gene expression reporters across successive growth cycles. Normalized relative reporter intensity from the first cycle was divided by that of the second cycle. a.u., arbitrary units.

Intrigued by this observation, we examined the spatial patterns of biofilm regrowth at the scale of a single biofilm. High-resolution imaging revealed that upon nutrient replenishment, cells in the PDBC grew to fill in regions previously occupied by cells in the founding biofilm (Fig. 5B, C). Comparing non-dispersed regions of individual biofilms across successive dispersal cycles revealed that regions which did not disperse in the initial cycle tended to also be non-dispersive in the subsequent dispersal event (Fig. 5C, Movie S7). These results suggest that heterogeneous matrix properties may render certain biofilm regions more amenable to dispersal than others, and that this spatial predisposition is maintained across multiple cycles of biofilm growth and dispersal.

Given that PDBC cells are already encased within a biofilm matrix, we next asked whether de novo gene expression of polysaccharide and protein components occurred during biofilm regeneration. To evaluate this possibility, we constructed a fluorescent promoter fusion to monitor *vps* matrix biosynthesis (*P_vpsL_-dL5*) along with our existing *rbmA* reporter (*P_rbmA_-dL5*), and quantified reporter activity over two cycles of biofilm formation and dispersal (Fig. 5D, E). Expression of *vpsL* and *rbmA* was markedly decreased during PDBC-initiated biofilm regrowth (∼8-fold and ∼17-fold) respectively, relative to initial biofilm development (Fig. 5F). Together, these findings suggest that PDBC cells may exploit existing matrix components to regenerate biofilms, enabling rapid re-establishment of surface communities while minimizing the energetic cost of rebuilding the matrix from scratch.

## Discussion

The natural world comprises diverse ecological niches, selecting for organisms adapted to variable environments. Microbes like *V. cholerae* persist across a range of niches by encoding phenotypic plasticity, enabling their transition between habitats as disparate as brackish water and the human intestine. Here, we show that genetically identical *V. cholerae* biofilm cells differentiate into two phenotypically distinct subpopulations during dispersal: while most cells exit the biofilms and return to the planktonic state, ∼10% of cells remain biofilm-associated and encased in the biofilm matrix. Further, we find that dispersed and PDBC cell populations differ primarily in their metabolic specialization indicating that dispersal-associated differentiation operates through mechanisms distinct from classical biofilm and regulatory pathways. We reason that biofilm dispersal is therefore not a uniform culminating step in the biofilm lifecycle but is instead consistent with a bet-hedging strategy in which the larger dispersal population optimizes chances of survival and niche relocation for this comparatively vulnerable population. Given the widespread reliance on biofilms for persistence and transmission, we hypothesize that differentiation during biofilm dispersal represents a general strategy for balancing dissemination and long-term survival with the opportunity for regrowth across diverse microbial systems.

We find that non-dispersed cells in the *V. cholerae* PDBC remain encased in the biofilm matrix which prevents the entry of competitors into the community, shields against environmental stresses, and enables long-term surface colonization. In natural environments, where competition for resources is ubiquitous, excluding competitors allows resident cells to monopolize available surfaces and the resources they provide. When fresh nutrients are provided, cells in the PDBC can re-initiate biofilm growth to repopulate an environmental niche, a process that is repeatable over multiple cycles. Interestingly, we find that PDBC cells do not substantially express biofilm matrix genes during regrowth, suggesting that they can use surrounding matrix as a scaffold to repopulate biofilm communities. We reason that this strategy enables cells repopulating the biofilm to reap the benefits of the biofilm lifestyle without the energy commitment involved in matrix production during the initial founding of a biofilm. Further investigation is required to determine how the existing matrix is remodeled during this process, or if, under certain circumstances, the existing matrix components could serve as an energy source for retained cells. In the context of infection, PDBCs could facilitate chronic infection where a reservoir of antibiotic tolerant cells can repopulate the host niche when challenges wane. Thus, an important area for future work will be to determine whether analogous post-dispersal biofilm-associated cell states exist in other bacterial pathogens and environmental microbes.

The maintenance of structural integrity in post-dispersal biofilms led us to the *V. cholerae* biofilm scaffolding protein RbmA, which we show is necessary and sufficient to retain cells in the PDBC. Given its role in altering matrix fluidity and cell packing in a biofilm, we propose that RbmA establishes heterogenous matrix properties across the biofilm, making regions differentially prone to dispersal (*35*, *43*). However, the regulatory mechanism(s) that predispose cells to dispersed and non-dispersed fates remain to be defined. Phenotypic diversification could be driven by stochastic changes in gene expression, or by cellular changes in response to environmental gradients like nutrients and oxygen that exist within structured communities (*44*, *45*). Varying growth conditions during biofilm development may establish cell fates either at dispersal, or earlier during development, possibly through the formation of distinct cell lineages better adapted to biofilm microenvironments. Notably, previous work has shown that biofilm dispersal in *V. cholerae* is a combination of position-dependent and random dispersal patterns (*26*), suggesting that multiple inputs may contribute to defining the fate of biofilm cells. Elucidating the molecular and physiological mechanisms that drive cell-cell variability and cell fate establishment in developing biofilms is an area of future work.

The canonical biofilm lifecycle progresses through well-defined stages beginning with irreversible surface attachment by a single planktonic cell, followed by cell division and the production of biofilm matrix components resulting in a mature multicellular community. The cycle culminates in dispersal where cells digest the surrounding matrix, return to the planktonic state and can seed a fresh biofilm elsewhere, closing the cycle. Here, we show that dispersal is not a uniform fate for biofilm cells - most cells disperse but they leave behind a PDBC harboring cells in a distinct physiological state that can re-enter the biofilm lifecycle and re-differentiate when provided favorable growth conditions. While phenotypic differentiation is a known feature of biofilm growth (*11*, *46*), we show here that dispersal itself becomes a cell fate decision, producing two cell populations, each capable of developing into biofilm communities, reframing dispersal from a uniform endpoint to a branching cell fate decision in the biofilm lifecycle.

## Acknowledgements

We thank the members of the Bridges Lab for their insightful comments and feedback on this manuscript. We thank Dr. Rich Olson for the Bap1-GFP fusion protein reagent used to visualize VPS in biofilms. We thank Dr. Robert van de Weerd for his advice and contributions to implementing the FAP imaging system.

## Author Contributions

Conceptualization: SK, AAB

Data Curation: SK, JAP, AAB

Formal analysis: SK, JAP, AAB

Funding acquisition: AAB

Investigation: SK, JAP, AAB

Methodology: SK, JAP, AAB

Project Administration: SK, AAB

Resources: SK, JAP, AAB

Supervision: AAB

Writing – original draft: SK, AAB

Writing – review & editing: SK, JAP, AAB

## Data, code, and materials availability

The source data used to generate all main and supporting figures in this work are available on Figshare. Biological materials used in this study are available upon request from Carnegie Mellon University. All custom processing scripts are available on Github.

## Materials and Methods

### Bacterial strains, growth conditions and fluorogenic reagents

The *Vibrio cholerae* parent strain used in this study was O1 El Tor biotype C6706str2. For propagation and cloning, strains were grown in lysogeny broth (LB) in liquid culture with shaking, or in petri plates supplemented with 1.5% agar at 30 °C. For biofilm growth in all experimental measurements, *V*. *cholerae* cells were grown in M9 medium containing glucose and casamino acids (1x M9 salts, 100 μM CaCl_2_, 2 mM MgSO_4_, 0.5% dextrose, 0.5% casamino acids). All strains used in this study are reported in Table S3. Antibiotics were used at the following concentrations, unless otherwise stated: kanamycin, 50 μg/mL; spectinomycin, 200 μg/mL; streptomycin, 400 μg/mL; chloramphenicol, 2 μg/mL; ampicillin, 100 μg/mL.

Fluorogenic compounds used in this study were acquired from the Carnegie Mellon University Technology Transfer & Enterprise Creation office. Fluorogen-activating proteins (FAPs) are single-chain variable fragment antibodies that bind to and restrict the rotation of small molecule fluorogens, resulting in amplification of fluorescence signal. The fluorogens used in this study were Malachite Green-2P (MG-2P – [λ_ex_ = 640 nm, λ_em_ = 680 nm]) and Thiazole Orange 1-2P (TO1-2P – [λ_ex_ = 510 nm, λ_em_ = 530 nm]). MG-2P and TO1-2P interact with their cognate FAPs, dL5 (referred to in the text as dL5-red) and AM2-2, and were excited using 640 nm and 488 nm laser lines, respectively. FAPs were expressed as translational fusions to the secretion signal of maltose binding protein to drive periplasmic localization (*26*, *47*). Fluorogens were dissolved and stored in ethanol with 1% acetic acid and were added to media at a final concentration of 1 µM (for MG-2P) or 1.5 µM (for TO1-2P) at the time of inoculation.

### Genetic manipulation and strain construction

Genetic manipulation of *V. cholerae* was performed by replacing genomic DNA with linear DNA introduced via natural transformation on chitin to stimulate competence (*48*, *49*). To generate linear DNA for transformation, DNA fragments were amplified from existing strains using PCR, fragments were stitched using overlap extension PCR as needed, or fragments were ordered as linear DNA from IDT. For constitutive labelling of cells with the fluorescent constructs *dL5-red* or *mNeonGreen,* expression was driven by the *tac* promoter expressed at the neutral locus, *vc1807*. In gene expression reporter strains, *AM2-2* was used for constitutive labeling, and expressed from a second neutral locus, *vc1378*, also driven by the *tac* promoter. To generate transcriptional fusions, DNA fragments immediately upstream of the start codon (300bp for *vpsL*, 500bp for *rbmA*) were stitched to *dL5-red* and expressed from *vc1807*. To construct the RbmA-3xFLAG fusion, a C-terminal 3xFLAG epitope was added to *rbmA* via a linker immediately upstream of the stop codon and introduced at the native *rbmA* locus. *P_bad_-rbmA-3xFLAG*, used for inducible expression, was introduced at the *vc1807* neutral locus. To generate strains with inducible *dL5-red*, a functional copy of *lacI,* which inhibits *P_tac_* activity, was expressed from the constitutive synthetic promoter PJ23106 at the *V. cholerae* native *lacIZ* locus. Expression of *dL5-red* was therefore repressed by LacI, and de-repressed by exogenous addition of 100 µM isopropyl β-D-1-thiogalactopyranoside (IPTG). Linear DNA oligonucleotides and synthetic linear DNA g-blocks were ordered from IDT and are reported in Table S4. In all cases, PCR and Sanger Sequencing (Genewiz) were used to verify genetic modifications.

### General biofilm growth and dispersal procedures

To perform microscopy of biofilm development and dispersal, initial overnight cultures were prepared by inoculating single *V. cholerae* colonies into 200 µL of LB media in polystyrene 96-well plates (Corning). Cultures were grown overnight at 30 °C with constant shaking under breathe-easier membranes (USA Scientific Inc.) to minimize evaporation. The next day, stationary-phase cultures were back-diluted to OD = 1 x 10^-5^ in 200 µL M9 media (see full recipe above) in polystyrene 96-well plates (Corning) for brightfield microscopy, or glass-bottom 96-well plates (Mattek) or 18-well chambered glass coverslips (Ibidi, #81817) for confocal microscopy, and incubated statically at 30 °C. This low seeding density stimulates *V. cholerae* biofilm formation, and spontaneous dispersal under these conditions enables the visualization of biofilm lifecycle dynamics of individual microcolony biofilms.(*25*)

### Spinning disc confocal microscopy

For spinning disk confocal microscopy, cells were grown in M9 media supplemented with the appropriate fluorogen(s) from the time of inoculation, and imaging was initiated 9 hours post-inoculation to capture dispersal, unless otherwise stated. Samples were imaged at 20-min intervals for 12 hours, until dispersal was complete. Cultures were maintained at 30 °C during imaging using a temperature-controlled chamber for microscopy (Oko Labs). Timelapse imaging was conducted on a motorized Nikon Ti2-E microscope equipped with a CREST X-Light V3 spinning disk unit and a back-thinned sCMOS camera (Hamamatsu Orca Fusion BT). Imaging utilized either a 100x silicone immersion objective (Nikon Plan Apochromat, NA 1.35), or a 20x air objective (Nikon Plan Apochromat, NA 0.75). Image acquisition was controlled by Nikon Elements software (Version 5.42.02), and illumination provided by an LDI-7 Laser Diode Illuminator (89-North).

To image PDBCs, cells were incubated until the completion of dispersal, dispersed cells were collected by pipetting, and PDBCs were washed three times with 1x PBS to remove any remaining dispersed cells and stained prior to imaging. To label dead cells, PDBCs were incubated with 1 µM of nucleic acid strain SYTOX Green (ThermoFisher, #S7020) in 1x PBS for 15 min and imaged at 488 nm. To visualize Vibrio polysaccharide (VPS), cells were probed with a fusion protein of Bap1 lacking the domain fused to GFP, a gift from Rich Olson (*36*, *50*). Samples were incubated in 100 µL M9 containing 1 µM of the Bap1-GFP fusion protein for 1 hour at room temperature and excited at 488 nm for microscopy. To capture biofilms at 48 hours post-inoculation to show persistence of the PDBC (Fig. S1A), imaging was performed at 10-hour, 24-hour, and 48-hour time points. For visualization of RbmA, cells expressing *rbmA-3xFLAG* were grown in media supplemented with a monoclonal Anti-FLAG-Cy3 Antibody (Sigma, #A9594) at a 1:1000 dilution from the time of inoculation. Samples were excited using the 561 nm laser every 30 min for 12 hours, starting at 7 hours post-inoculation.

### Three-dimensional biofilm analysis

To analyze all spinning-disc biofilm timelapses captured at high resolution (using a 100x objective), images were first denoised using the Denoise.ai module available in Nikon Elements software (Version 5.42.02) and a rolling ball background subtraction (radius = 50) was performed in FIJI (Version 2.16.0) (*51*). To correct for x-y drift between timepoints, timelapse images were registered using a Fourier phase-correlation approach with subpixel refinement as described by Guizar-Sicairos and colleagues (*52*). 3D stacks of individual time points were segmented using the cellpose-SAM (Version 4)(*53*) algorithm using 2500 iterations with a smoothing factor of 1.0 and minimum cell diameter of 20 voxels. Biofilm cells were differentiated from planktonic cells via two criteria based on cell tracks generated using the Trackpy package (Version 0.6.4)(*54*) in Python (Version 3.10.11): First, to exclude transient interactions between planktonic cells and the biofilm, only cells present in the timelapse for more than a defined number of timepoints were considered part of the biofilm. Second, new cells at every timepoint were only considered part of a biofilm if their centroids were within a defined distance from the biofilm at the previous timepoint, with the largest connected component of the biofilm mask at the first timepoint serving as the initial biofilm configuration. The stringency of this distance threshold was increased starting with timepoints close to the onset of biofilm dispersal. The number of biofilm-associated cells per time frame was obtained and used to assess the percentage of biofilm cells over the course of dispersal.

For analysis of RbmA-3xFLAG localization, pre-processing and segmentation were the same as described above. To obtain RbmA-3xFLAG fluorescence intensity over time, we first defined the biofilm region at each time point by generating a 2D convex hull of the maximum projections of image stacks that had been isotropically rescaled. Projections were then combined to generate a global biofilm mask which was applied to every slice of the rescaled z-stack. Using this approach, we calculated total raw fluorescence intensity within the hull as well as the total number of cells at each time point. To generate kymographs (Fig. 3F), the same biofilm masks generated above were used to define biofilm regions. Images were downsampled 2-fold for analysis. Radial distances were computed relative to the geometric centroid (forced to the bottom slice) of the global hull, and mean fluorescence was calculated within concentric radial bins of approximately 2 microns. Fluorescence image analysis were performed using the Julia programming language (Version 1.11.2) (*55*).

### Microscopy of transcriptional reporters in biofilms

For assessment of fluorescent gene expression reporters, timelapse spinning-disc microscopy was performed at 1-hour intervals for 16 hours using a 20x objective to capture initial biofilm growth and dispersal. We found that reporter output was too low to enable higher resolution timelapse imaging without photobleaching. To assess reporter output in the second cycle of biofilm formation, 24-hour old PDBCs were washed three times with 1x PBS and exchanged into fresh M9 media supplemented with fluorogens. For biofilm regrowth, which was captured using identical microscopy settings, a new field of view was imaged to minimize the effects of prolonged imaging on cell viability and photobleaching.

To quantify whole-biofilm fluorescence intensity of promoter fusions from confocal images, sum intensity projections over z-slices (∼20 µm depth) were generated for constitutive (*P_tac_-AM2-2*) and reporter channels (transcriptional fusions to *dL5-red*) and used for downstream analyses. For each channel, the mean intensity of the first timepoint was subtracted from all timepoints, followed by Gaussian blurring (σ = 2 pixels) to reduce noise. Masks of biofilm regions were obtained using intensity-based thresholding on the *P_tac_-AM2-2* constitutive channel. Resulting masks were applied to both channels independently to obtain the mean fluorescence intensity from entire biofilm regions for each time point. Reporter fluorescence was obtained by dividing fluorescence intensities from the reporter channel by that of the constitutive channel and each trace was normalized to its peak signal before plotting.

### Biofilm invasion

Cells expressing dL5-red and mNeonGreen were seeded separately to form biofilms in chambered glass coverslips. After dispersal, dL5-red-labeled dispersed cells were removed, and PDBCs were washed three times with 1x PBS. mNeonGreen*-*expressing dispersed cells were transferred onto dL5-red-expressing PDBCs in M9 media containing fluorogen. Samples were imaged for 3 hours at 15-min intervals by confocal microscopy using a 100x objective.

### *dL5-red* induction in PDBCs

To assess the propensity of dispersed and PDBC cells to produce RNA and protein after dispersal, the two cell populations were separated but maintained in identical conditioned media. To collect conditioned media, biofilms were grown in 6-well plates until the completion of dispersal (16 hours post-inoculation), culture fluids were aspirated and cells were removed by 10 min of centrifugation at 3,200 x *g*, followed by filtration through a sterile 0.2 µm filter. In separate biofilm cultures, cells harboring a functional copy of *lacI* and *P_tac_*-*dL5-red* (which is repressed by LacI) were used to seed initial biofilm formation. After the completion of biofilm dispersal (16 hours post-inoculation), dispersed cells were collected and diluted 1:20 into conditioned media (dilution determined empirically, see Competition Experiments for details). Simultaneously, PDBCs were washed three times with conditioned media to remove any remaining dispersed cells. Both dispersed and PDBC populations were then separately exposed to 100 µM IPTG to induce *dL5-red* expression. Samples were incubated in an Agilent Biospa robotic incubator and transferred to an Agilent Biotek Cytation 1 plate reader microscope to measure bulk fluorescence intensity at 30-min intervals after induction using a Cy5 filter cube (Ex/Em 620/680 nm).

### Brightfield Imaging Assays

To capture biofilm formation and dispersal (Fig. 2B, Fig. 5A) samples were imaged using low magnification brightfield timelapse microscopy, as described previously (*25*, *56*). To induce *rbmA* expression, arabinose was added at indicated concentrations at the time of biofilm inoculation. To capture biofilm regrowth following dispersal, dispersed cells were collected and discarded every 24 hours, cells in the PDBC were washed three times with 1x PBS and subsequently provided fresh M9 media. Samples were incubated in an Agilent Biospa robotic incubator. At 1-hour intervals, microplates were transferred to an Agilent Biotek Cytation 1 plate reader microscope, and imaged using a 10x (Olympus Plan Fluorite, NA 0.3, air) objective. Instruments were controlled using the Biospa OnDemand software (Version 1.04) and Biotek Gen5 (Version 3.12). Biofilm quantification was based on the light attenuation by verticalized multicellular communities. To segment biofilm structures, pixel values were inverted, local contrast was normalized, images were blurred using a Gaussian filter and were segmented based on a linear intensity threshold. Resulting binary masks were applied to raw images to obtain the total amount of attenuated light within a given field of view as a measure of biofilm biomass.(*25*, *56*)

### Competition experiments

To perform competitions between dispersed and PDBC cell populations, strains expressing either mNeonGreen or dL5-red were initially grown separately as biofilm cultures in 96-well polystyrene plates for 16 hours, until the completion of dispersal. Dispersed cells were subsequently collected by pipetting, subjected to 16,000 x *g* for 1 min in microcentrifuge tubes, and resuspended in fresh media. From the same initial samples, PDBCs were washed three times with 1x PBS to remove remaining dispersed cells. Competitions were performed by mixing equal numbers of dispersed cells and PDBC cells expressing distinct fluorescent constructs, in the presence or absence of environmental stressors.

To achieve equal starting cell numbers, we empirically determined the appropriate dilution of dispersed cells by measuring bulk cellular fluorescence from each population of a strain constitutively expressing *dL5-red*. Fluorescence intensities across dilutions of dispersed cells were compared to those of dL5-red-labeled PDBCs. This approach was validated across multiple days and biological replicates to account for experimental variability. Colony-forming units were not used as a quantification metric due to the difficulty of liberating single cells from established PDBCs.

For competition experiments, dL5-red–labeled PDBCs (which retain fluorescence in post-dispersal biofilms) were mixed with mNeonGreen-labeled dispersed cells. In parallel, dispersed cells expressing both fluorescent markers were competed against one another to confirm that any observed differences in growth were attributable to lifestyle rather than fluorescent labeling. To expose cell populations to abiotic threats, we first determined sub-MIC concentrations of abiotic stressors in dispersed and non-dispersed populations in monoculture (125 µM hydrogen peroxide and 3.1 µM of chloramphenicol). For exposure to lytic phage S5, cells were competed at a multiplicity of infection of 0.1 in phage medium (LB containing 0.5% dextrose and 10 mM CaCl_2_) instead of M9.

Survival of each population was measured using bulk fluorescence measurements on an Agilent Biotek Cytation 5 plate reader driven by Biotek Gen5 (Version 3.12) software. For each condition, monochromators were tuned to measure culture fluorescence for dL5-red (Ex/Em 635/680 nm) and mNeonGreen (Ex/Em 495/525 nm). In each experiment, planktonic monocultures were grown in parallel to generate an independent linear calibration curve correlating optical density (OD_600_) to fluorescence intensity for each channel, which were used to convert fluorescence values into a computed optical density. Only exponential-phase growth data were used for these measurements as we found that fluorescence signal for both reporters continued to accumulate in stationary phase despite reduced cell division. For each strain, population growth was calculated as the fold change in computed optical density from the initial time point to the final time point (10 hours post competition initiation). Population survival was quantified as the ratio of fold change in treated samples relative to untreated controls for each fluorescently labeled strain. Statistical analysis was performed on R. Schematics and figures were made on Inkscape (Version 1.3.2).

### RNA Isolation and Transcriptomic Analysis

For RNA isolation, triplicate overnight cultures of *V. cholerae* were diluted to OD = 1 x 10^-5^ into M9 media in 96-well polystyrene plates to stimulate static biofilm formation at 30 °C. After the completion of dispersal, at 16 hours post-inoculation, dispersed cells were collected by centrifugation at 16,000 x *g* for 3 min in microcentrifuge tubes and resuspended in RNAprotect (NEB) and RNA Lysis buffer (NEB). PDBCs were washed three times with 1x PBS to remove any dispersed cells and incubated in RNAprotect (NEB) and RNA Lysis buffer (NEB) for 15 min at room temperature. RNA was isolated using the NEB RNA Isolation Kit following manufacturer’s instructions. The concentration and purity of isolated RNA was measured on a Nanodrop (Thermo). Samples were frozen at -80 °C and shipped on dry ice to SeqCenter (https://www.seqcenter.com/rna-sequencing/) for sequencing. 12 million paired end reads sequencing and intermediate analysis package were selected for each sample.

Library preparation was performed using Illumina’s Stranded Total RNA Prep Ligation with Ribo-Zero Plus kit and 10bp unique dual indices (UDI). Sequencing was performed on a NovaSeq X Plus, producing paired end 150bp reads. Demultiplexing, quality control, and adapter trimming was performed with bcl-convert (Version 4.1.5). Read mapping was performed using HISAT2 and read quantification was performed using Subreads’s FeatureCounts functionality (*57*, *58*). Read counts were normalized using edgeR’s TMM (Trimmed Mean of M values) algorithm, and differential analysis performed using edgeR’s glmQLFTest (*58*, *59*). Pathway Overrepresentation Analysis was performed in R using ClusterProfiler with all significantly differentially expressed genes (Log_2_fold > 1 and *P*-value < 0.05) (*60*). Gene Set enrichment Analysis was performed using the fgsea library with defined gene lists as input, given in Table S2 (*61*). All plots were generated using ggplot2.

### c-di-GMP Quantification

To quantify c-di-GMP in dispersed and non-dispersed cell populations, wild-type *V. cholerae* was grown as biofilms in 96-well polystyrene plates until the completion of dispersal (16 hours post-inoculation), at which point the two populations were separated. PDBCs were washed three times using 1x PBS. Cell lysis was performed by exchange into 1x BugBuster Protein Extraction Reagent (Millipore Sigma, #70584) (100 µL/well) followed by incubation at 50 °C for 90 min. Samples were then centrifuged in microcentrifuge tubes at 16,000 x *g* for 5 min to remove cell debris, and the resulting supernatants were used for downstream processing.

Sample c-di-GMP concentrations were measured using a Cyclic-di-GMP ELISA Kit (Cayman Chemical, #501780) following manufacturer’s instructions. Quantification is based on a competitive assay where sample c-di-GMP displaces a fluorescent analog, resulting in an inverse correlation between fluorescence signal and sample c-di-GMP concentration. Absorbance was measured at 450 nm using the Agilent Biotek Cytation 5 plate reader driven by Biotek Gen5 (Version 3.12). Absolute concentrations in each sample were obtained using a standard curve and then normalized to sample protein concentration (A_280_) measured using the Nanodrop (Thermo) to account for differential lysis. For each experiment, we included control samples treated with polyamines norspermidine and spermidine at 100 µM, which have established roles in elevating and reducing intracellular c-di-GMP levels respectively (*62*).

## Supplementary Figures

**Figure S1:**
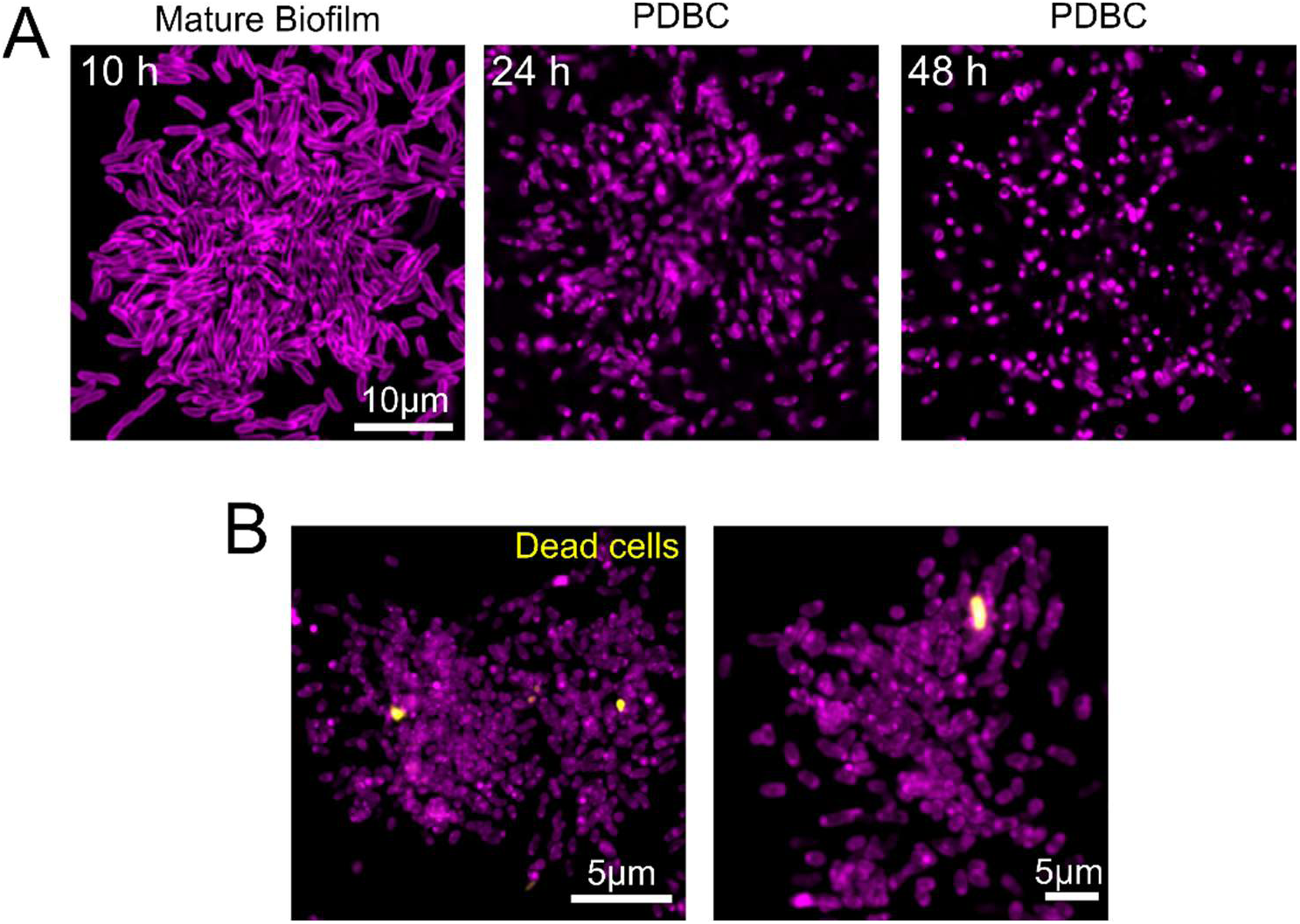
Cells in *V. cholerae* PDBCs are alive and retained for extended periods of time. (A) Confocal micrographs (bottom slice) of cells imaged at the indicated time points post biofilm inoculation. Cell shape changes during starvation result in the observed cell rounding in late-stage PDBCs. (B) Representative sum projections of replicate *V. cholerae* PDBCs stained with the nucleic acid stain SYTOX Green, *N* = 3 biological replicates. Magenta represents all cells constitutively expressing *dL5-red*, and yellow represents SYTOX signal.

**Figure S2:**
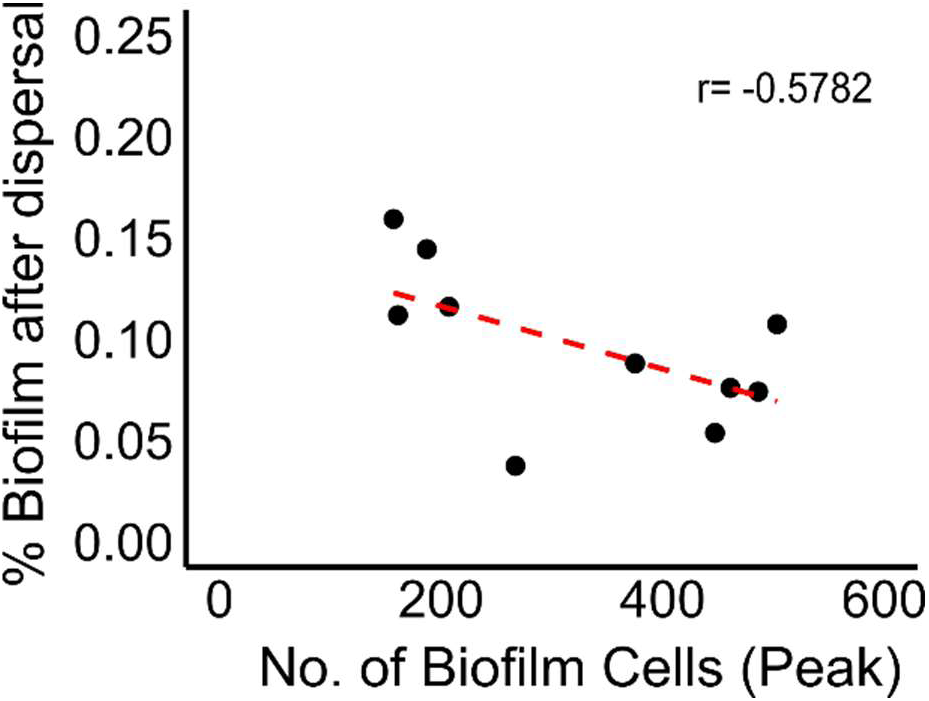
Correlation between the size of a mature biofilm and the proportion of cells remaining after dispersal. Each point represents a single biofilm and red dashed line shows the trendline of *N* = 10 replicate biofilms. Correlation coefficient (r) of -0.57 indicates a modest negative correlation.

**Figure S3:**
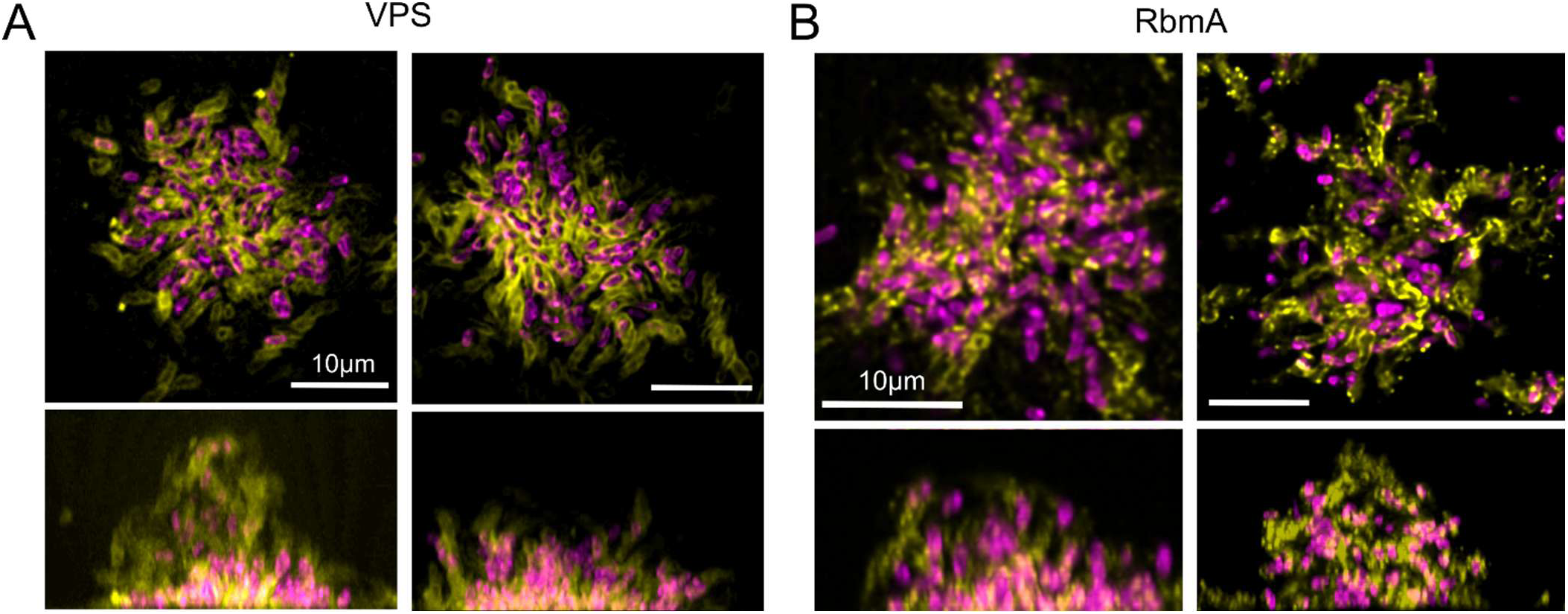
Replicate *V. cholerae* PDBCs probed for matrix components. (A) Representative confocal micrographs (bottom slice and x-z projections), of PDBCs probed for VPS. (B) Representative confocal micrographs (bottom slice and x-z projections) of PDBCs probed for RbmA-3xFLAG. Magenta represents cells constitutively expressing *dL5-red*, yellow represents indicated matrix components. *N* = 3 biological replicates for both imaging conditions.

**Figure S4:**
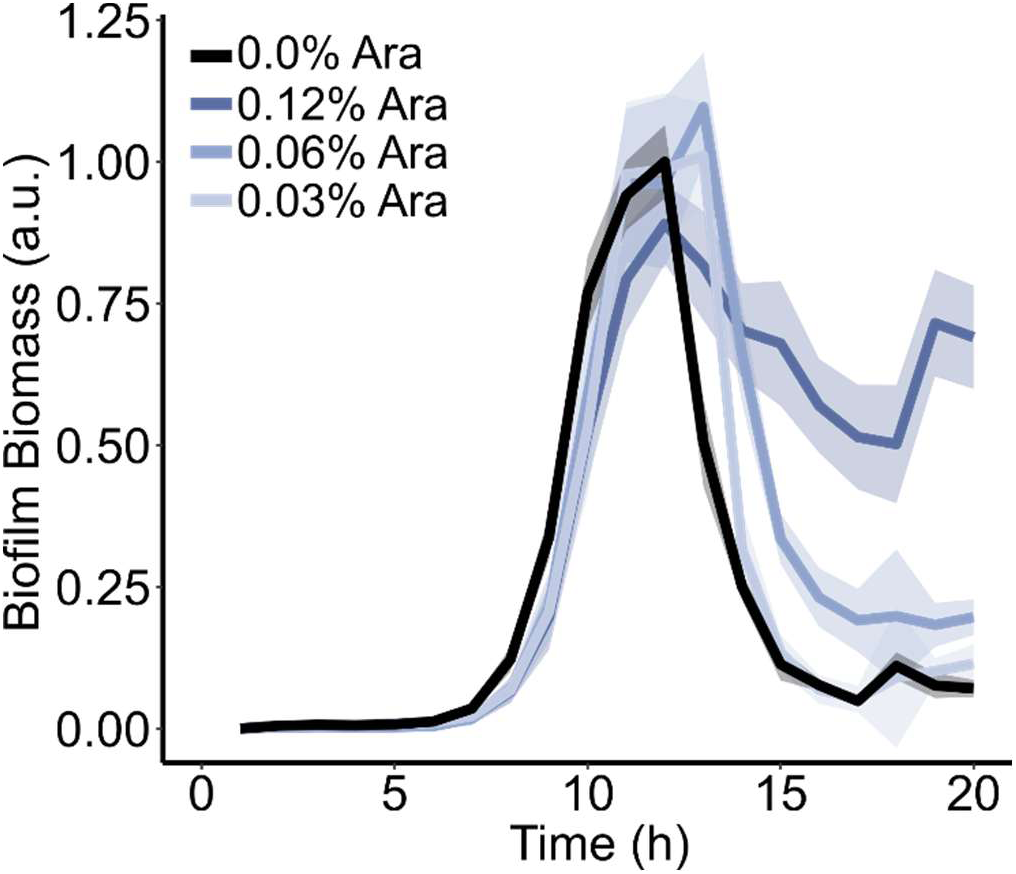
Effect of *rbmA* expression on biofilm biomass over time. Quantification of biofilm formation over time of a Δ*rbmA* strain induced with the indicated concentrations of arabinose to induce *rbmA* expression. Lines represent means of *N* = 3 biological and 3 technical replicates per condition ± SD (shaded).

**Figure S5:**
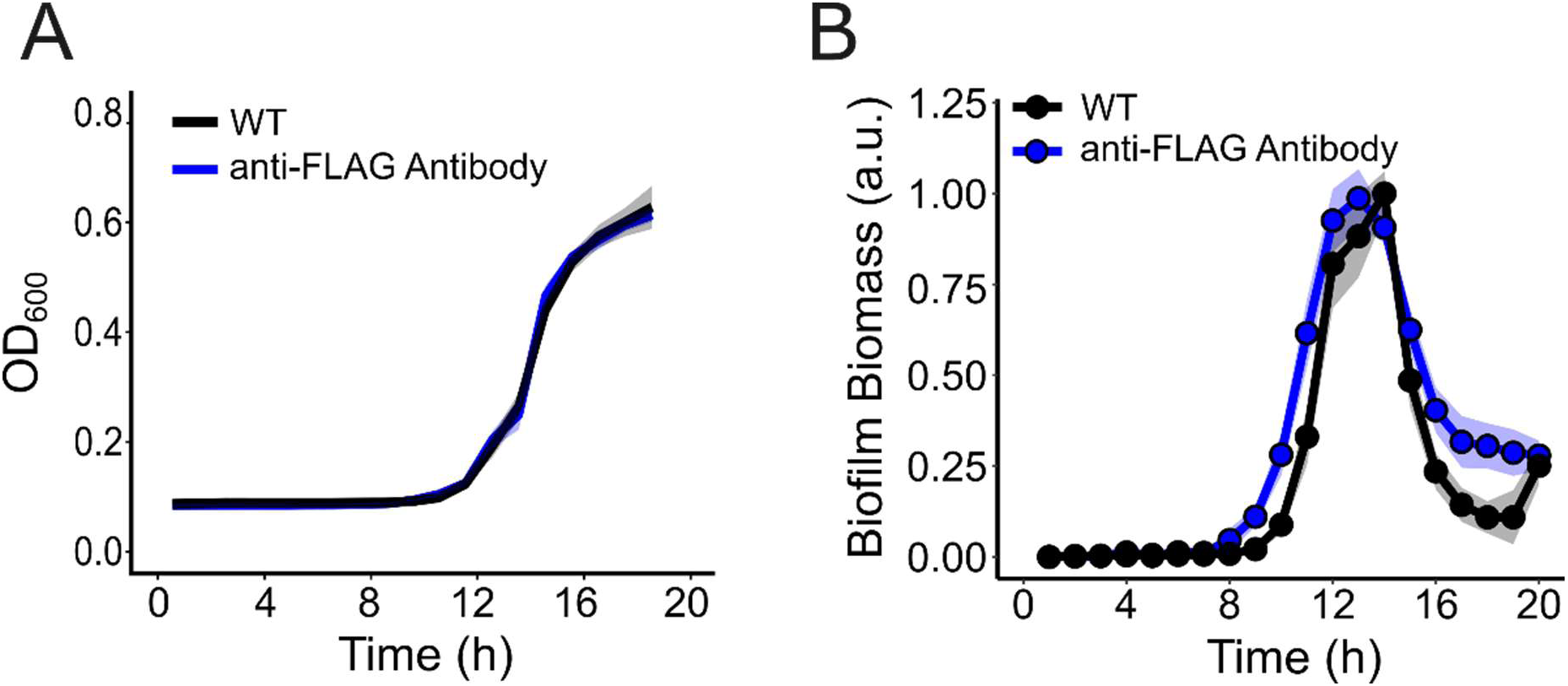
Growth of wild-type *V. cholerae* in the presence of anti-FLAG antibody. (A) Population growth quantified using optical density (OD_600_) measurements of cells on their own, or in the presence of a 1:1000 dilution of anti-FLAG antibody. Lines represent means of *N* = 3 biological and 3 technical replicates of each condition, ± SD (shaded). (B) Quantification of biofilm formation over time of cells grown on their own, or in the presence of anti-FLAG antibody. Points represent means of *N* = 3 biological and 3 technical replicates of each strain, ± SD (shaded). a.u., arbitrary units.

**Figure S6:**
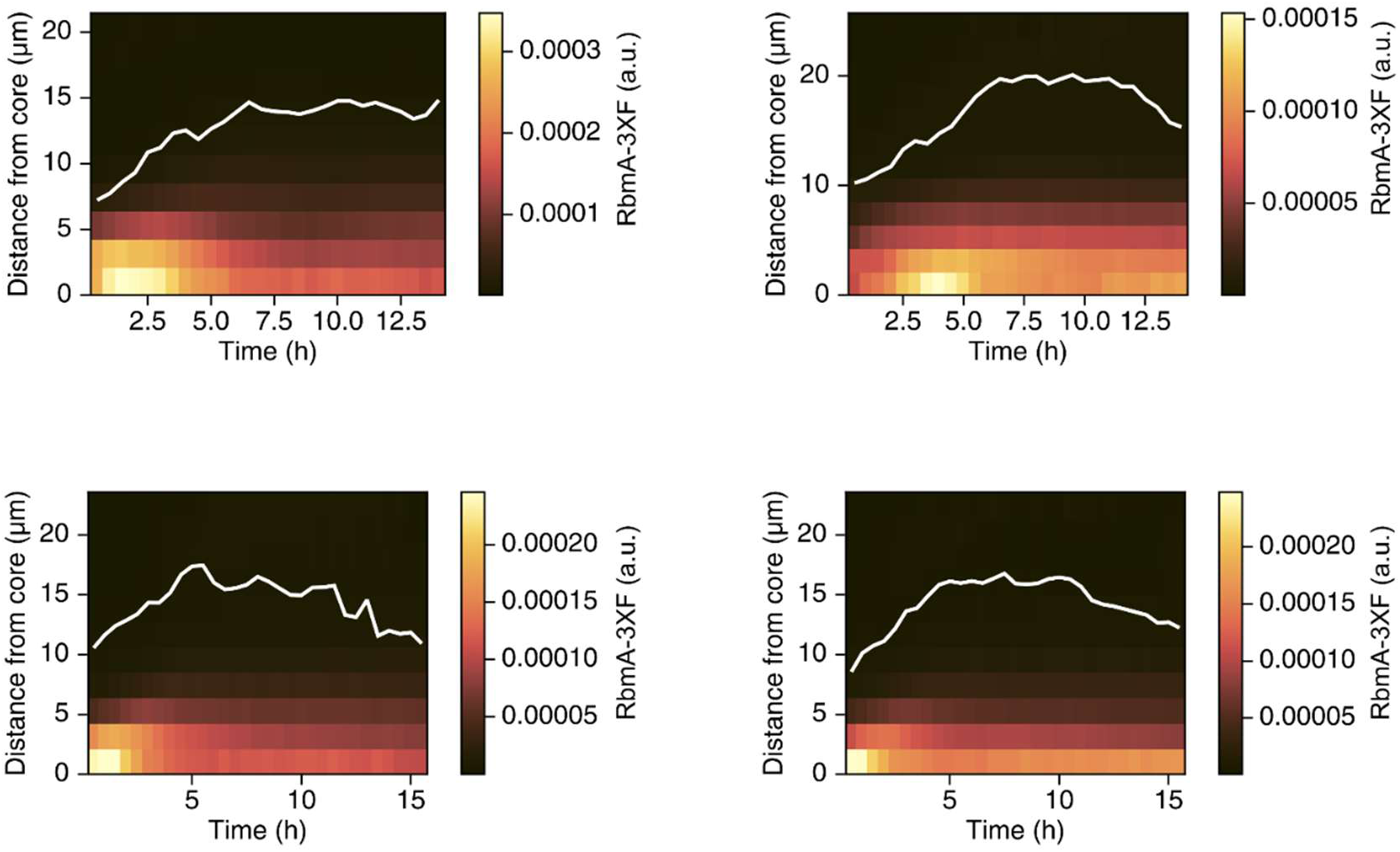
Spatiotemporal pattern kymographs of RbmA-3xFLAG fluorescence in individual replicates. Representative kymographs of individual biofilms, quantifying total RbmA-3xFLAG fluorescence intensity over time in radial bins at the indicated distances from the biofilm core. White line represents biofilm boundary. a.u., arbitrary units.

**Figure S7:**
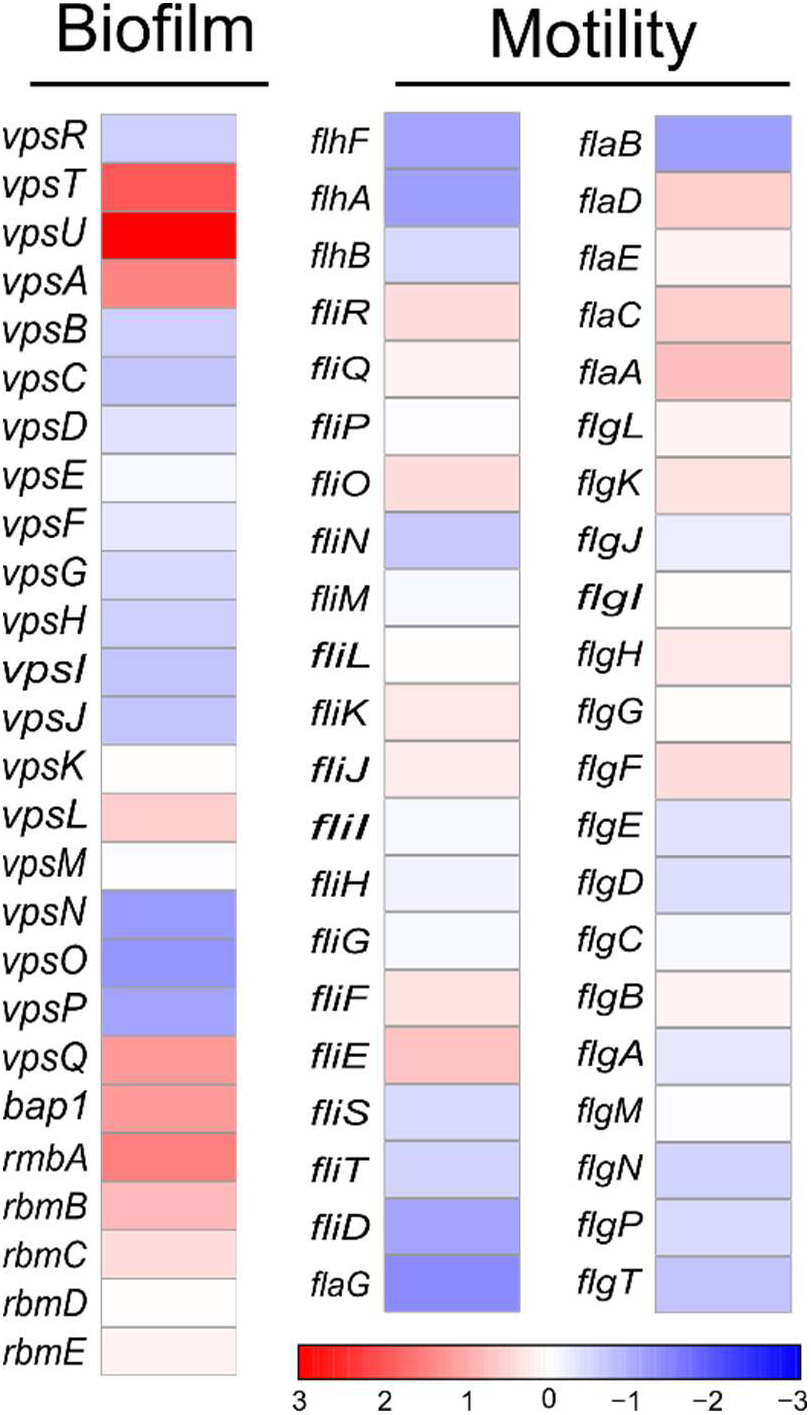
Gene expression changes of known biofilm and motility genes in PDBCs relative to dispersed cells. Heatmap depicting Log_2_ Fold Changes of known biofilm and motility genes in PDBCs relative to dispersed cells.

**Figure S8:**
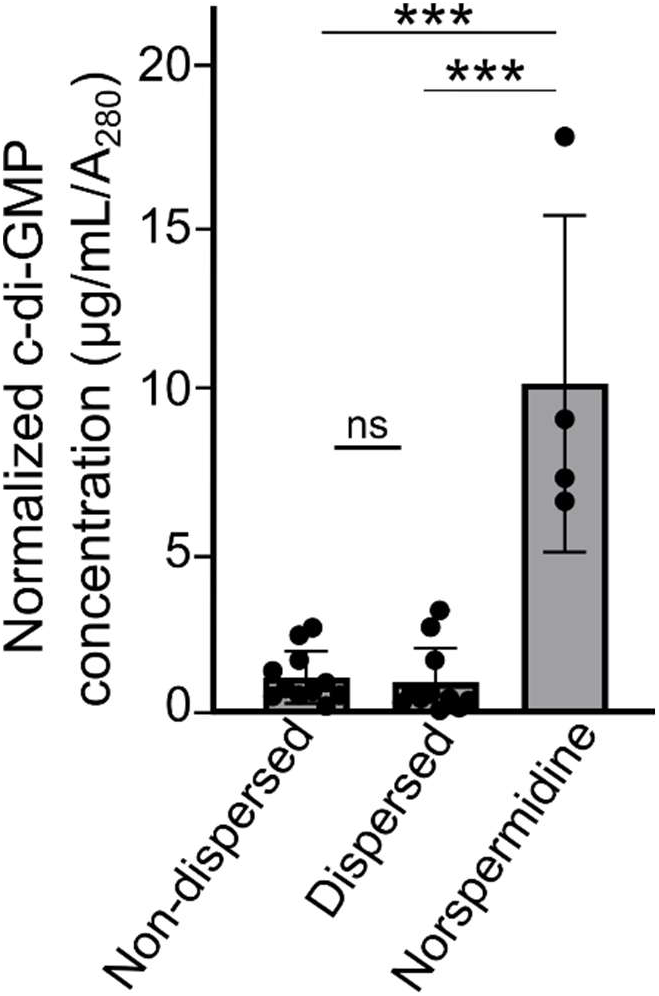
c-di-GMP in dispersed and non-dispersed cell populations. Total c-di-GMP concentrations in dispersed and non-dispersed cell populations normalized to sample protein concentration (A_280_), compared to treatment with 100 µM norspermidine. Unpaired *t*-tests were used for statistical analysis, ****, *P* < 0.0001; ns, *P* > 0.05. Dispersed and PDBC data represent *N* = 3 biological and 3 technical replicates. Norspermidine data represents *N* = 2 biological and 2 technical replicates.

**Figure S9:**
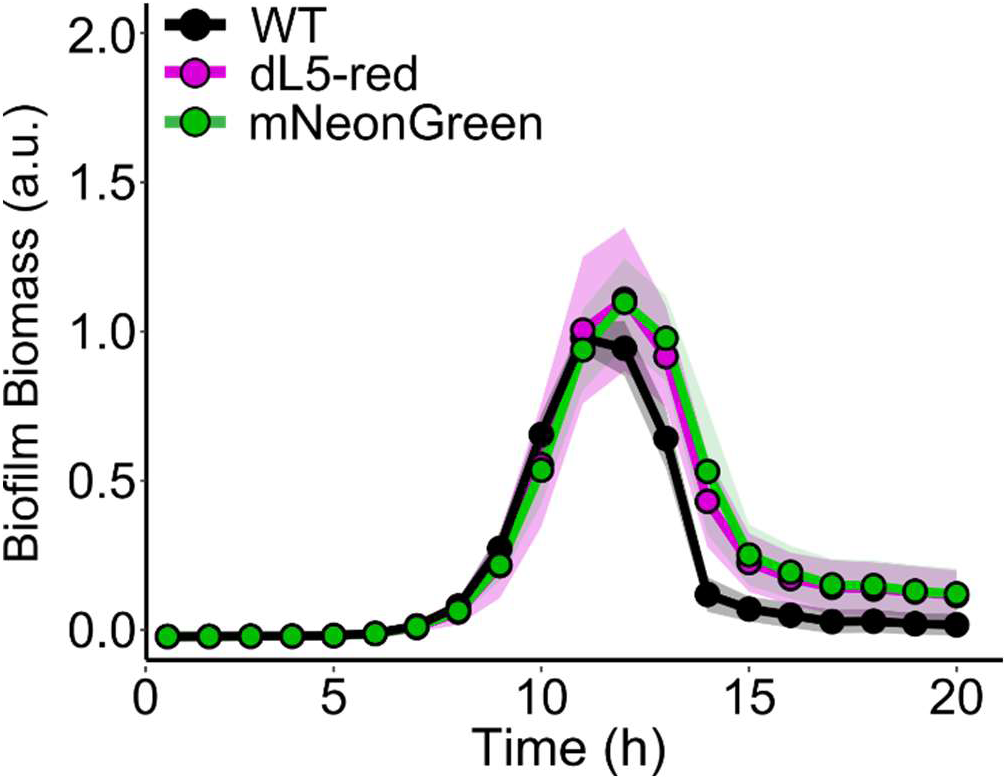
Isogenic strains used for competition experiments undergo comparable dispersal. Quantification of biofilm formation in strains used for competition experiments relative to the wild-type strain. Lines represent means of *N* = 3 biological and 3 technical replicates per strain, ± SD (shaded). a.u., arbitrary units.

## Supplementary Tables

**Table S1:** Significantly differentially regulated genes in non-dispersed cells relative to dispersed cells. Added as a separate Excel file.

**Table S2:** Input Gene Sets and Statistics for Gene Set Enrichment Analyses.

| Pathway | P-val | p-adj | ES | NES | Input Gene List | Enriched Genes |
| --- | --- | --- | --- | --- | --- | --- |
| Biofilm | 0.601881 | 0.874916 | 0.33608 | 0.889685 | VC_0916; VC_0917; VC_0918;<br>VC_0919; VC_0920; VC_0921;<br>VC_0922; VC_0923; VC_0924;<br>VC_0925; VC_0926; VC_0927;<br>VC_0934; VC_0935; VC_0936;<br>VC_0937; VC_0938; VC_0939 | VC_0916; VC0917;<br>VC_0939 |
| Flagella | 0.729097 | 0.874916 | -0.24591 | -0.8451 | VC_2120; VC_2121; VC_2122;<br>VC_2123; VC_2124; VC_2125;<br>VC_2126; VC_2127; VC_2128;<br>VC_2129; VC_2130; VC_2131;<br>VC_2132; VC_2133; VC_2134;<br>VC_2138; VC_2139; VC_2140;<br>VC_2141; VC_2142; VC_2143;<br>VC_2144; VC_2187; VC_2188;<br>VC_2190; VC_2191; VC_2192;<br>VC_2193; VC_2194; VC_2195;<br>VC_2196; VC_2197; VC_2198;<br>VC_2199; VC_2200 | VC_2141; VC_2142;<br>VC_2140; VC_2125;<br>VC_2139; VC_2120;<br>VC_2138; VC_2198;<br>VC_2197; VC_2192;<br>VC_2131; VC_2126;<br>VC_2130; VC_2132;<br>VC_2199; VC_2123;<br>VC_2193; VC_2195;<br>VC_2127; VC_2122;<br>VC_2200; VC_2190;<br>VC_2144; VC_2129;<br>VC_2194; VC_2128;<br>VC_2133; VC_2191;<br>VC_2196; VC_2121;<br>VC_2124; VC_2143;<br>VC_2187; VC_2134;<br>VC_2188 |
| Stringent Response | 0.523891 | 0.877637 | 0.456451 | 0.973626 | VC_2451; VC_1224; VC_2710;<br>VC_0437; VC_0596; VC_0534;<br>VC_0767; VC_2277 | VC_2277; VC_0596;<br>VC_0437; VC_0767;<br>VC_2710 |

p-val, $P$ -value; p-adj, adjusted $P$ -value; ES, Enrichment Score; NES, Normalized Enrichment Score.

**Table S3:** Strain List.

| Strain No. | Genotype | Antibiotic Resistance | Origin |
| --- | --- | --- | --- |
| AB_Vc_707 | *Vibrio cholerae* C6706str2 | - | Bassler Lab |
| AB_Vc_761 | $\Delta vc1807::Cm^R$ | $Cm^R$ | NT of AB_Vc_707 |
| RW_Vc_1892 | $\Delta vc1807::Ptac$-SS(MBP)-DL5::$Spec^R$ | $Spec^R$ | Prentice et al., 2024 |
| SK_Vc_2266 | $\Delta vc1807::Ptac$-SS(MBP)-DL5::$Kan^R$ | $Kan^R$ | NT of AB_Vc_707 |
| AB_Vc_310 | $\Delta vc1807::Ptac$-mNeonGreen(co)-$Spec^R$ | $Spec^R$ | NT of AB_Vc_707 |
| JP_Vc_1427 | $\Delta rbmA \Delta 1807::Kan^R$ | $Kan^R$ | Prentice et al., 2024 |
| JP_Vc_1940 | $\Delta rbmA \Delta 1807::Spec^R$ | $Spec^R$ | NT of JP_Vc_1427 |
| JP_Vc_1970 | $\Delta rbmA Ptac$-ss(MBP)-dL5 $\Delta vc1807::Spec^R$ | $Spec^R$ | Prentice et al., 2024 |
| SK_Vc_2289 | $rbmA$-3xF $\Delta vc1807::Cm^R$ | $Cm^R$ | NT of JP_Vc_1940 |
| SK_Vc_2292 | $rbmA$-3xF $\Delta vc1807::Ptac$-SS(MBP)-DL5::$Spec^R$ | $Spec^R$ | NT of SK_Vc_2289 |
| SK_Vc_2112 | $\Delta rbmA \Delta vc1807::Pbad$-$rbmA$-3xF::$Kan^R$ | $Kan^R$ | NT of JP_Vc_1940 |
| SK_Vc_2283 | $\Delta vc1807::PrbmA$-SS(MBP)-DL5::$Spec^R$ | $Spec^R$ | NT of AB_Vc_707 |
| AB_Vc_2124 | $\Delta vc1807::PvpsL$-ssMBP-DL5::$Kan^R$ | $Kan^R$ | NT of AB_Vc_707 |
| SK_Vc_2287 | $\Delta vc1807::PrbmA$-SS(MBP)-DL5::$Spec^R \Delta vc1378::Ptac$-ssMBP-AM2-2::$Amp^R$ | $Amp^R$<br>$Spec^R$ | NT of AB_Vc_2283 |
| AB_Vc_2131 | $\Delta vc1807::PvpsL$-ssMBP-DL5::$Kan^R \Delta vc1378::Ptac$-ssMBP-AM2-2::$Amp^R$ | $Amp^R$ $Kan^R$ | NT of AB_Vc_2124 |
| SK_Vc_2142 | $\Delta vc1807::Ptac$-SS(MBP)-DL5::$Spec^R \Delta lacIZ::PJ23106_{lacI}::Kan^R$ | $Kan^R$ $Spec^R$ | NT of RW_Vc_1892 |
| SK_Vc_2136 | $\Delta vpsU \Delta vc1807::Kan^R$ | $Kan^R$ | NT of AB_Vc_707 |
| SK_Vc_2180 | $\Delta vpsU \Delta vc1807::Ptac$-SS(MBP)-DL5::$Spec^R$ | $Spec^R$ | NT of SK_Vc_2136 |

**Table S4:**
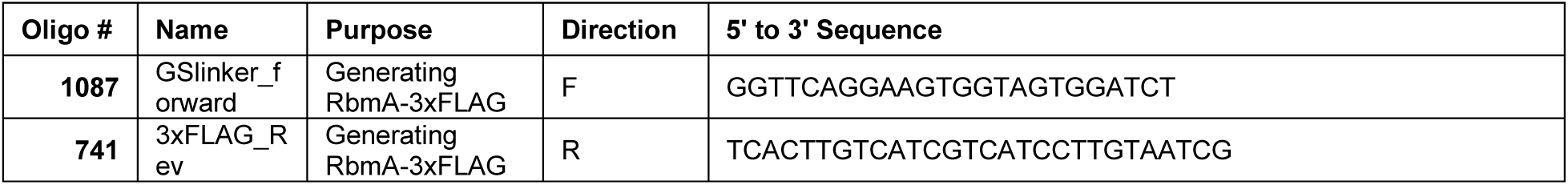
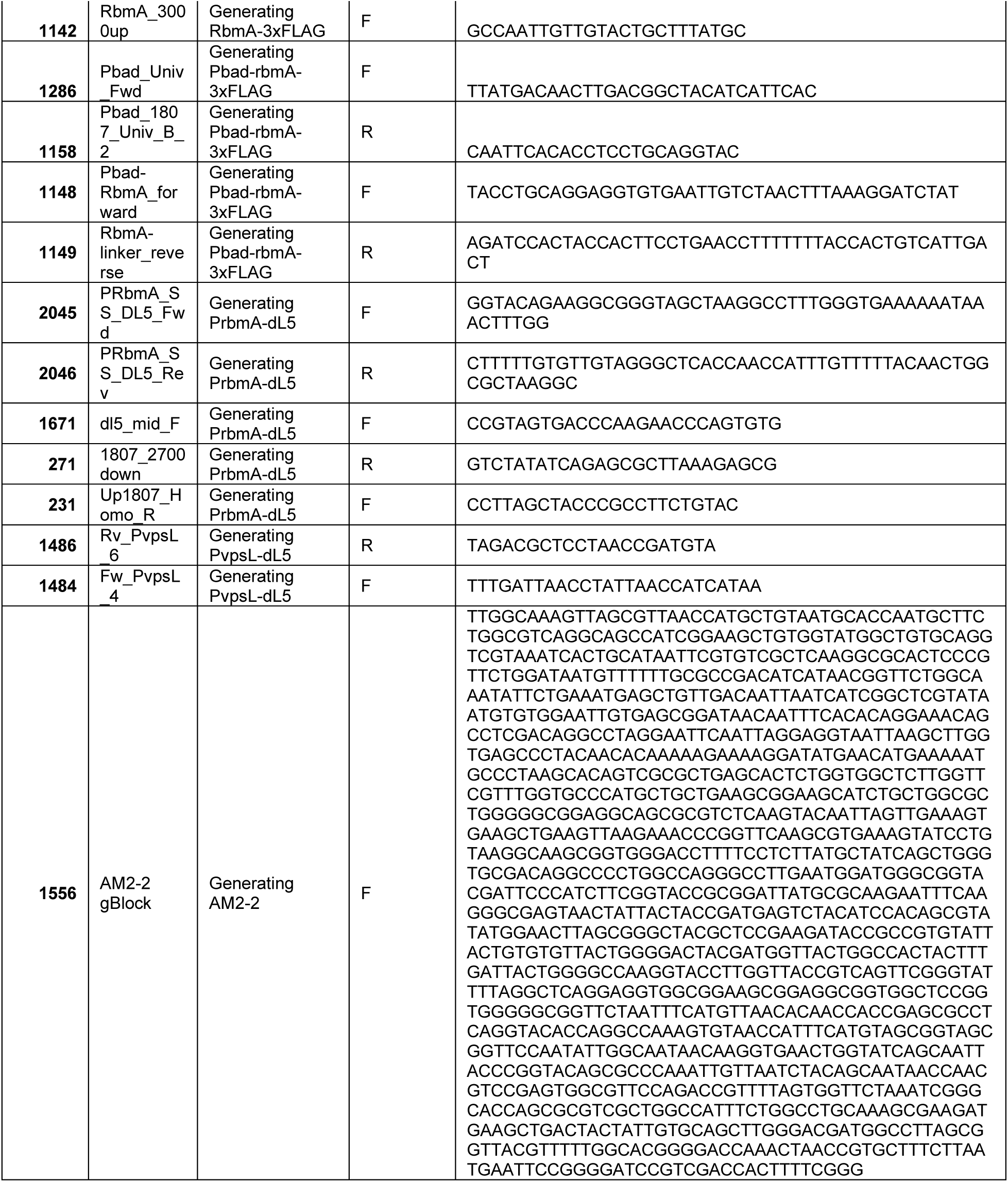
Oligonucleotides used in this study.

## SUPPLEMENTARY MOVIES

**Movie S1: Representative timelapse of WT *V. cholerae* biofilm dispersal.** Single x-y slice (left panel) and x-z projection (right panel). Scale bar and time stamp as indicated.

**Movie S2: Representative two-color imaging experiment.** Single x-y slice (left panel) and x-z projection (right panel). Magenta represents PDBC cells, green represents dispersed cells, imaged immediately after cultures were mixed. Scale bar and time stamp as indicated.

**Movie S3: Representative timelapse of Δ*rbmA V. cholerae* biofilm dispersal.** Single x-y slice (left panel) and x-z projection (right panel). Scale bar and time stamp as indicated.

**Movie S4: Representative timelapse of WT *V. cholerae* biofilm development probed for RbmA-3xFLAG.** Single x-y slice (left panel) and x-z projection (right panel). Magena represents cells, and yellow represents RbmA-3xFLAG probed with an anti-FLAG antibody. Scale bar and time stamp as indicated.

**Movie S5: Representative timelapse of *V. cholerae* PDBCs following dL5-red induction.** Single x-y slice imaged immediately after the addition of IPTG. Scale bar and time stamp as indicated.

**Movie S6: Representative timelapse of WT *V. cholerae* biofilm growth and dispersal over 100 hours.** Low magnification brightfield images of WT *V. cholerae* over four cycles of biofilm growth and dispersal. Scale bar and time stamp as indicated.

**Movie S7: Representative timelapse of a single WT *V. cholerae* biofilm growth and dispersal 48 hours.** Single x-y slice. Scale bar and time stamp as indicated.

